# Orc6 at replication fork enables efficient mismatch repair

**DOI:** 10.1101/2021.05.10.443400

**Authors:** Yo-Chuen Lin, Arindam Chakraborty, Dazhen Liu, Jaba Mitra, Lyudmila Y Kadyrova, Rosaline Y.C. Hsu, Mariam K. Arif, Sneha Adusumilli, Taekjip Ha, Farid A Kadyrov, Kannanganattu V. Prasanth, Supriya G. Prasanth

## Abstract

In eukaryotes, the Origin Recognition Complex (ORC) is required for the initiation of DNA replication. The smallest subunit of ORC, Orc6, is essential for pre-replication complex (pre-RC) assembly and cell viability in yeast and for cytokinesis in metazoans. However, unlike other ORC components, the role of human Orc6 in replication remains to be resolved. Here, we identify an unexpected role for hOrc6, which is to promote S-phase progression post pre-RC assembly and DNA damage response. Orc6 localizes at the replication fork and is an accessory factor of the mismatch repair (MMR) complex. In response to oxidative damage during S-phase, often repaired by MMR, Orc6 facilitates MMR complex assembly and activity, without which the checkpoint signaling is abrogated. Mechanistically, Orc6 directly binds to MutSα and enhances the chromatin-association of MutLα, thus enabling efficient mismatch repair. Based on this, we conclude that hOrc6 plays a fundamental role in genome surveillance during S-phase.

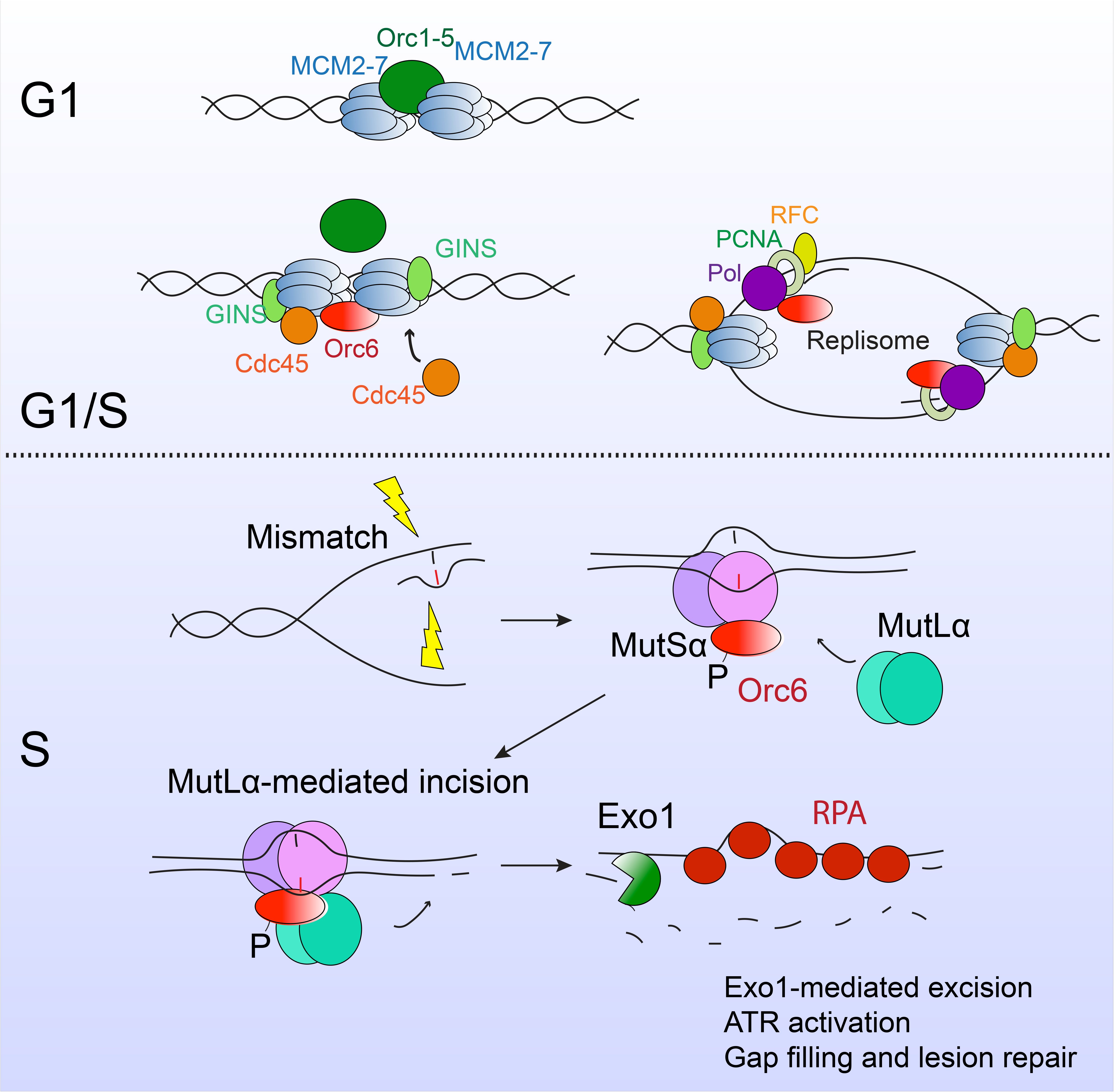

**Highlights:**

- Human Orc6 is dispensable for G1 licensing, but required for S-phase progression
- Human Orc6 at the replication fork is an accessory factor for MMR complex
- Depletion of hOrc6 sensitizes cells to DNA damage and impairs ATR activation
- Human Orc6 regulates MMR complex assembly and activity

## Introduction

Accurate duplication of the genetic material and faithful transmission of genomic information are critical to maintain genome stability. Errors in DNA replication and repair mechanisms are deleterious and cause genetic aberrations leading to malignant cellular transformation and tumorigenesis. Origin recognition complex (ORC) proteins are critical for the initiation of DNA replication (Bell and Stillman, 1992), and the individual subunits of ORC also play vital roles in several non-preRC functions, including heterochromatin organization, telomere maintenance, centrosome duplication and cytokinesis (Chesnokov, 2007; Sasaki and Gilbert, 2007). Mutations within several *ORC* genes, including *ORC1, ORC4 and ORC6*, have also been linked to Meier-Gorlin Syndrome, a rare genetic disorder in children characterized by primordial dwarfism (Bicknell et al., 2011; Bleichert et al., 2013; Guernsey et al., 2011; Hossain and Stillman, 2012; Kuo et al., 2012).

ORC serves as the landing pad for the assembly of the multiprotein pre-replication complex at the origins of replication during G1 (Bell and Dutta, 2002). The smallest subunit of ORC, Orc6, is highly dynamic with respect to its association with the other ORC components (Dhar and Dutta, 2000; Siddiqui and Stillman, 2007; Vashee et al., 2001). Orc6 is an integral part of ORC in yeast and *Drosophila* (Bell and Stillman, 1992; Chesnokov et al., 2001), but only weakly associates with ORC in human and *Xenopus* (Dhar et al., 2001; Dhar and Dutta, 2000; Gillespie et al., 2001; Vashee et al., 2001). Orc6 possesses DNA binding ability and is believed to be critical for DNA replication initiation in all eukaryotes (Balasov et al., 2007; Chen et al., 2007; Li et al., 2018; Semple et al., 2006; Thomae et al., 2008); however, human Orc6 and Orc1-5 can bind to DNA independently (Thomae et al., 2011). In human cells, Orc6 is present in sub-stoichiometric levels to the other ORC subunits. There is evidence that hOrc6 protein directly binds to the Orc3 subunit and integrates as a part of ORC *in vivo* in human cell lines (Siddiqui and Stillman, 2007). However, a significant fraction of hOrc6 is not associated with the ORC complex, suggesting that hOrc6 is involved in ORC-independent functions within the cell (Dhar et al., 2001; Vashee et al., 2001). In support of this, metazoan Orc6 is also required for cytokinesis and this function of Orc6 is facilitated by its binding to septin proteins (Balasov et al., 2007, 2009; Chesnokov et al., 2003; Prasanth et al., 2002). In yeast, Orc6 is dispensable for progression through mitosis and cytokinesis, but the depletion of Orc6 after pre-RC assembly has been shown to impair replication origin firing (Semple et al., 2006). However, no such role for Orc6 has been evaluated in higher eukaryotes.

Accurate duplication of the genetic material and correction of errors during S phase are efficiently coordinated with the machinery that ensures genomic integrity (Cook, 2009; Cortez, 2019). Mismatches occurring during DNA replication are recognized and removed by the Mismatch Repair (MMR) system. The key components of the MMR system that have been identified to date are MutSα (MSH2-MSH6 complex), MutSβ (MSH2-MSH3 complex), MutLα (MLH1-PMS2 complex), RFC-loaded PCNA, and Exonuclease 1 (EXO1) (Kunkel and Erie, 2015; Modrich, 2006). The association of MMR machinery to the replication fork is facilitated by PCNA, an essential replication factor that is also required for MMR (Flores-Rozas et al., 2000; Gu et al., 1998; Hombauer et al., 2011; Kadyrov et al., 2006; Kleczkowska et al., 2001; Umar et al., 1996). In eukaryotes, MMR is active throughout the cell cycle with the highest activity observed during S phase (Edelbrock et al., 2009; Schmidt and Hombauer, 2016; Schroering et al., 2007). Several replication accessory factors, chromatin associated factors and epigenetic regulators also influence MMR (Awwad and Ayoub, 2015; Kadyrova et al., 2011; Loughery et al., 2011; Schopf et al., 2012; Yuan et al., 2004), yet the functional relevance of these interactions has not been well understood. It is also worth noting that mutations within several replication factors result in defective MMR, however, the mechanistic studies are lacking.

We report that hOrc6 associates with the replication fork and is primarily required for DNA replication progression but not for G1 licensing. During S-phase, Orc6 forms an integral component of the mismatch repair (MMR) complex, and controls MMR complex assembly and activity. Further, in response to oxidative damage, often repaired by MMR, Orc6 gets phosphorylated during S-phase, which in turn inhibits DNA replication progression. Loss of Orc6 results in defective MMR activity, resulting in loss of ATR signaling. Based on our results, we conclude that hOrc6 has a fundamental role in genome surveillance during S-phase.

## RESULTS

### Orc6 is a component of the replication fork

Orc6 is known to display robust DNA binding ability in metazoans (Balasov et al., 2007; Xu et al., 2020). In order to understand the type of DNA structure that facilitates the binding of hOrc6, we performed Single Molecule Pull down (SiMPull) assays (Jain et al., 2011). We observed that hOrc6 bound more tightly to replication fork structures (Kd 6.34+0.49nM) compared to ssDNA (19.24+5nM) and dsDNA (15.82+1.24nM) structures (Figure S1A and S1B). In order to test if hOrc6 associates with the replication fork *in vivo*, we performed isolation of Proteins On Nascent DNA (iPOND) (Sirbu et al., 2013). Nascent DNA was labeled with EdU, conjugated with biotin, and proteins associated with biotin-EdU-labeled DNA were pulled down. Thymidine chase experiment was used as the mature chromatin control. Orc6 showed accumulation on nascent DNA, whereas it did not enrich on the mature DNA (Figure 1A). To confirm this data and gain quantifiable results, we performed quantitative in situ analysis of protein interactions at DNA replication forks (SIRF) (Roy et al., 2018). Nascent DNA was labeled with EdU, conjugated with biotin, then proximity ligation assay (PLA) was performed to determine the association between hOrc6 and biotin-EdU-labeled DNA. SIRF experiments also showed that hOrc6 associated with nascent DNA (Figure 1B). Similar to our observation, hOrc6 was found in proteomic screens of proteins enriched in nascent DNA by nascent chromatin capture (SILAC log_2_ ratio = 0.76+/-.45) (Alabert et al., 2014) as well as by iPOND (log_2_ ratio = 1.28) (Wessel et al., 2019), suggesting its association with replication forks in human cells. Furthermore, immunoprecipitation experiments (Figure 1C) and SiMPull (Figure S1C and S1D) demonstrated that hOrc6 interacted with several replication fork components, including the single-stranded DNA binding protein RPA, the DNA clamp PCNA, and the clamp loader RFC. These results demonstrate that hOrc6 is enriched at the replication fork and associates with the fork components, implying that Orc6 in human cells could be involved in functions downstream of pre-RC assembly.

**Figure 1.**
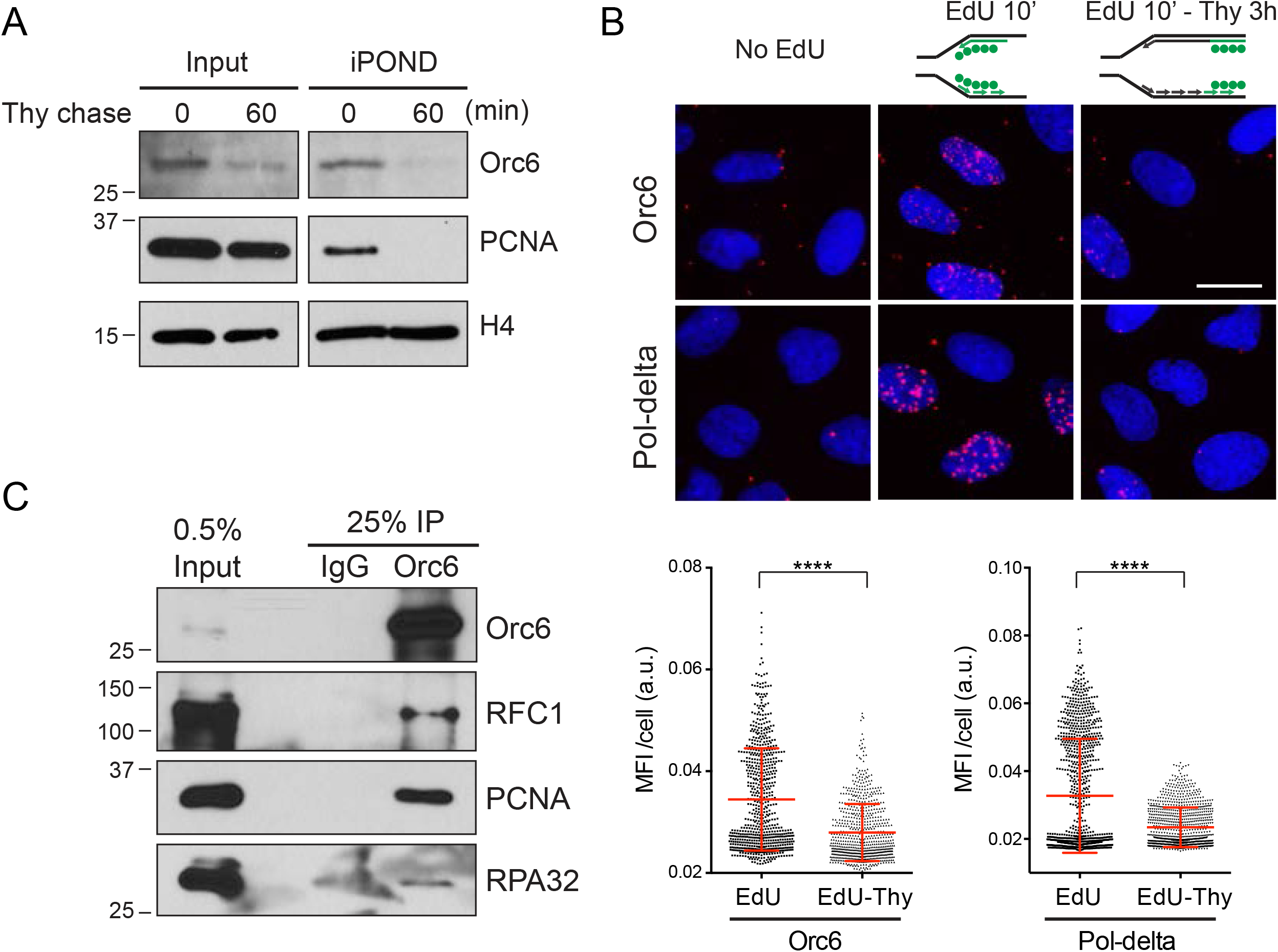
Orc6 is at replication fork and associates with fork components. (A) Western blot of iPOND. Thymidine chase 0 min for nascent DNA and 60 min for mature DNA. PCNA as a positive control for nascent DNA. (B) Upper panel: representative images for SIRF. Red foci indicate the association. PLA of Pol-delta/Edu-biotin served as a positive control. DAPI as counterstain. Scale bar, 25µm. Lower panel: quantification results. Experiments were performed in triplicates and one representative experiment is shown; n > 700 for each group. Mean ± SD. ****p < 0.0001 by unpaired two-tailed Student’s t test. MFI: mean fluorescence intensity. a.u.: arbitrary unit. (C) Immunoprecipitation of endogenous Orc6 from U2OS cells. See also Figure S1.

### Orc6 is required for accurate S phase progression

In yeast and *Drosophila*, Orc6 is involved in origin licensing as part of ORC (Bleichert et al., 2013; Miller et al., 2019); however, the function of human Orc6 has not been fully examined. The depletion of Orc6 in human cells with an intact p53-mediated checkpoint status caused a decrease in S phase population with a concomitant reduction in the chromatin association of PCNA (Figure 2A and S2A). The inability of the hOrc6-depleted cells to progress through S phase could either be due to reduced origin firing caused by defects in preRC assembly and/or inhibition of replication elongation. It has been shown that in metazoans the C-terminus of Orc6 interacts with Orc3 and hence recruits Orc6 to the core ORC (Bleichert et al., 2013). We addressed if the association of hOrc6 to ORC and the suggested role for ORC in licensing is responsible for the S-phase defects in Orc6-depleted cells. To test this, we performed complementation experiments using either full-length Orc6 or C-terminal truncated Orc6 (a.a. 1-187), which doesn’t associate with ORC. We observed that the C-terminal truncation mutant rescued the S-phase defect to a similar extent to that of the full length Orc6 (Figure S2B). This suggests that, in human cells, the S phase defect of Orc6 depletion might not be due to defects of ORC function in origin licensing. To evaluate this further, we used flow cytometry to measure MCM licensing status in conjugation with cell cycle analysis (Matson et al., 2017). Cells were extracted using detergent before fixing to remove soluble MCMs. The chromatin bound MCMs served as an indicator of origin licensing and were detected by immunostaining with MCM3 antibody (Figure 2B). Strikingly, hOrc6-depleted cells showed a similar level of chromatin bound MCMs in G1 population to control cells, suggesting that the depletion of Orc6 did not alter the MCM association to the chromatin (Figure 2C and 2D). In contrast, Orc1 knockdown showed a severe licensing defect as indicated by dramatic reduction of chromatin-bound MCMs. However, even with proper licensing, hOrc6-depleted cells were unable to efficiently enter S phase.

**Figure 2.**
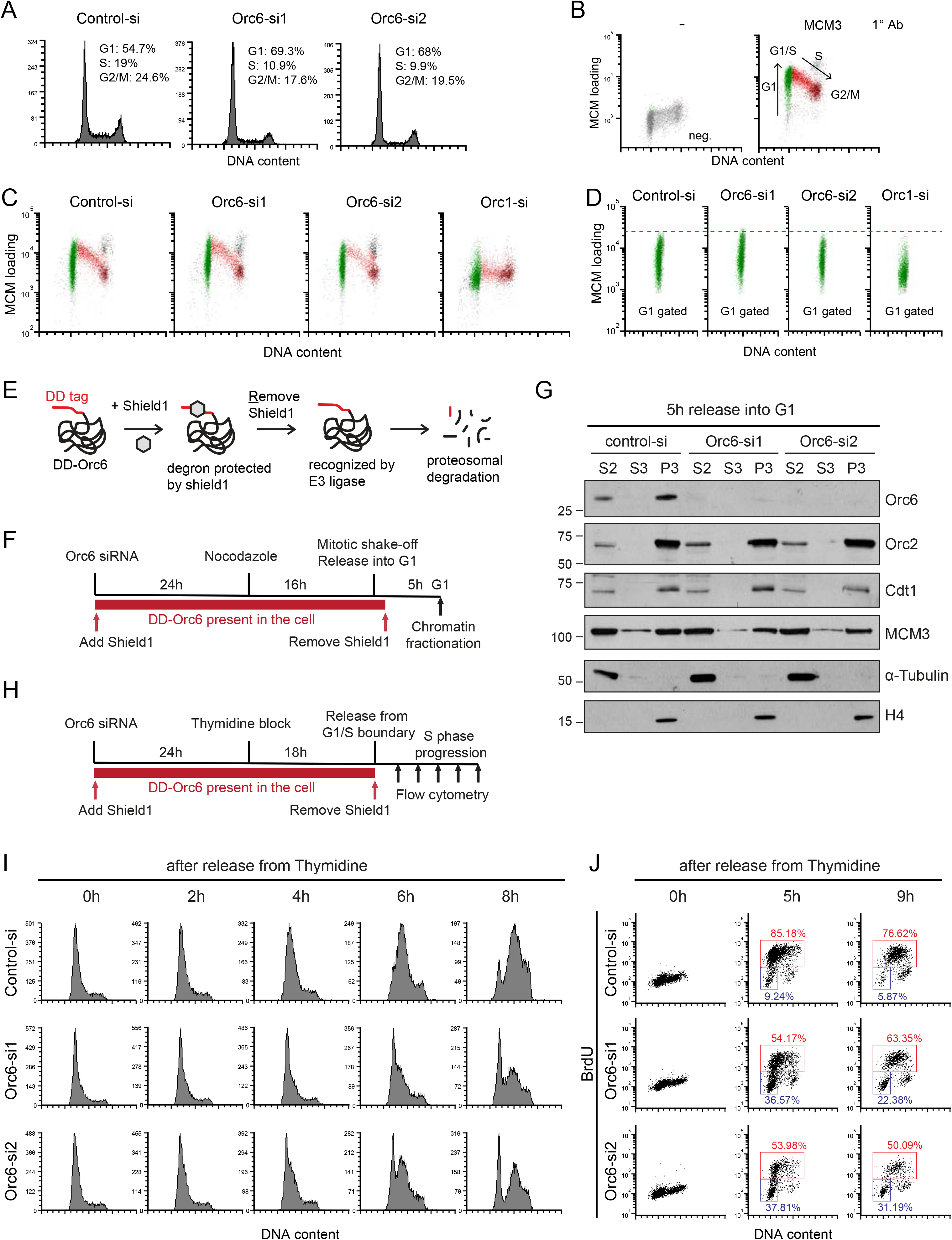
Orc6 is not essential for G1 licensing but required for proper S phase progression. (A) Cell cycle profile of Orc6 knockdown cells. (B) Flow cytometric analysis of asynchronized U2OS cells to measure chromatin bound MCM in conjugation with PI staining cell cycle profile. Left panel: MCM3 antibody omitted as negative gating for MCM staining. Right panel: illustration of cell cycle progression; loading of MCM on chromatin increases through G1 (green); S phase cells (red) with increasing DNA content and decreasing chromatin MCM, until reaching G2/M (maroon). (C) MCM-PI flow of Orc6 and Orc1 knockdown cells. (D) G1 populations from (C) were highlighted for comparing the MCM loading. (E) Schematic illustration of the DD degron system for controlling the degradation of DD-Orc6. (F) Schematic of the protocol for specifically depleting Orc6 in G1 phase. (G) Western blot analysis of the G1 phase chromatin fractionation. S2, cytosolic; S3, nuclear soluble; P3, chromatin fraction. α-tubulin and H4 serve as loading control for cytosolic and chromatin fraction, respectively. (H) Schematic of the protocol for specifically depleting Orc6 in S phase. (I) S phase progression determined by PI flow cytometry. (J) S phase progression determined by BrdU-PI flow cytometry. See also Figure S2.

In *Saccharomyces cerevisiae*, when Orc6 was depleted during late G1, MCM proteins were displaced from chromatin, and cells failed to progress through S phase, suggesting that efficiency of replication origin firing was compromised (Semple et al., 2006). However, similar role has not been evaluated for Orc6 in higher eukaryotes. To further assess if hOrc6 is truly dispensable for G1 licensing and functionally separate the role of hOrc6 in G1 from post-G1, we utilized a degron system, which allows degradation of Orc6 by ubiquitin-proteasomal degradation at any specific time point during the cell cycle (Giri et al., 2015). We tagged hOrc6 with a destruction domain (DD).

This DD tag is recognized by the proteasomal machinery and facilitates the rapid degradation of the DD-Orc6. In the presence of Shield1, a molecule that masks the DD tag, DD-Orc6 is prevented from degradation (Figure 2E). DD-Orc6 could substitute for endogenous Orc6, as the tagged-hOrc6 rescued the cell cycle defects observed in cells depleted of endogenous hOrc6 (Figure S2C and S2D; compare samples 2 and 3). We depleted the endogenous Orc6 using siRNA targeting the 3’-UTR of Orc6 in U2OS cells stably expressing DD-Orc6 and carried out the experiment in the presence of Shield1. We then synchronized the DD-Orc6-expressing cells depleted of endogenous Orc6 into early G1 phase (Figure 2F), and degraded DD-Orc6 by removing Shield1 from the medium. By determining the loading of preRC components onto chromatin in G1 cells lacking Orc6, we monitored the chromatin association of preRC components (Orc2, Cdt1, MCM3). The chromatin loading of these factors remain unaffected in the cells depleted of hOrc6, again suggesting that Orc6 is dispensable for replication licensing in human cells (Figure 2G).

Using the DD-Orc6 stably expressing cells, we evaluated the effect of hOrc6 loss in post-G1 cells on progression through S phase (Figure 2H). The degradation of hOrc6 in post-G1 cells resulted in slower progression through S phase. PI flow profile at 4-8 hrs time point post thymidine release revealed that more cells progressed through S phase in control cells compared to Orc6-depleted (Orc6-si1 & -si2 lacking endogenous as well as DD-Orc6) cells (Figure 2I). BrdU-PI profile further corroborated these results demonstrating the defects in S phase progression of Thymidine-released post-G1 cells in the absence of hOrc6 (Figure 2J).

In order to determine why the loss of hOrc6 in post-G1 cells caused defects in S phase progression, we determined the status of chromatin loading of preRC and pre-IC components in these cells (Figure 3A). We did not observe any defect in the total and chromatin-associated fraction of ORCs and MCMs in presence or absence of hOrc6 (Figure 3B). Similarly, PCNA and the DNA polymerases displayed comparable chromatin loading in the presence or absence of hOrc6 (Figure 3B). However, cells depleted of hOrc6 only during G1/S and S phase showed significant reduction in the chromatin association of Cdc45, a critical component of the CMG helicase (Figure 3B). The WT-Orc6 was able to efficiently rescue the Cdc45 loading in Orc6-depleted cells (Figure 3C). We further tested the role of hOrc6 in the chromatin loading of Cdc45 using a parallel cell biological approach. Using an *in vivo* reporter assay (Shen et al., 2010), we tethered MCM2 to a heterochromatic gene locus in U2OS 2-6-3 cells and examined the recruitment of endogenous Cdc45 to that site. In control cells, we found that the tethering of MCM was sufficient to recruit endogenous Cdc45 to the locus (52.8%; n = 36) (Figure 3D). However, MCM2 failed to recruit Cdc45 to the locus in cells depleted of endogenous hOrc6 (27.5%; n = 40) (Figure 3D). Both biochemical and cell biological approaches demonstrate that hOrc6 play vital roles in the recruitment of Cdc45 to the chromatin in post-G1 cells. All of these results imply that hOrc6 plays an essential role in DNA replication.

**Figure 3.**
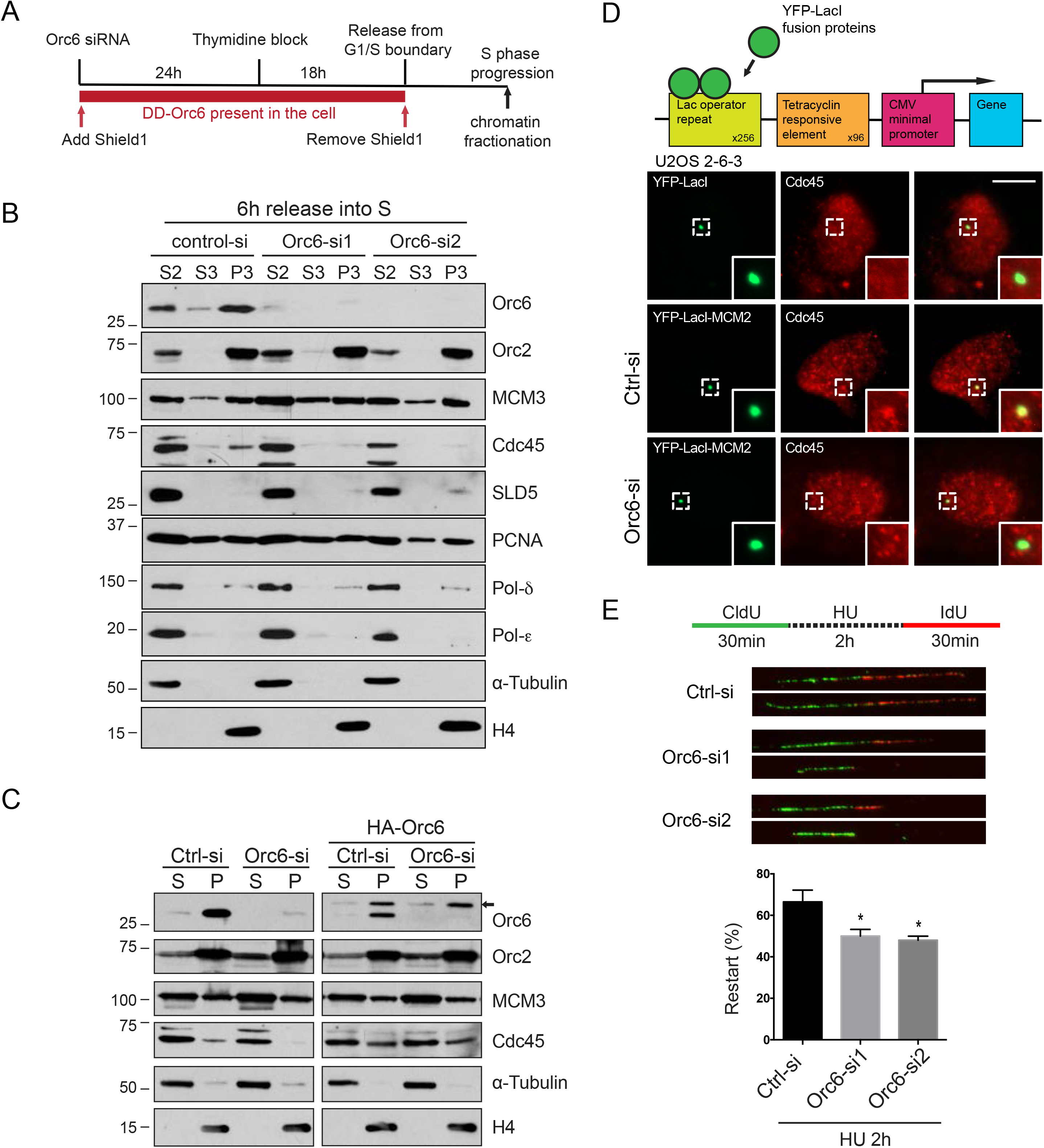
Depletion of Orc6 causes reduced cdc45 chromatin loading and defects in replication fork restart. (A) Schematic of the protocol for specifically depleting Orc6 in S phase. (B) Western blot analysis of the S phase chromatin fractionation. (C) Similar in (B), western blot showing the chromatin fractionation of Orc6-depleted samples expressing HA-Orc6. *Arrow* indicates HA-Orc6. (D) Upper panel: schematic of the heterochromatic locus stably integrated in U2OS 2-6-3 cells. LacI fusion proteins are forcibly tethered to the LacO repeats, and YFP is fused to LacI for visualizing the loci. Lower panel: images of cdc45 recruitment to the heterochromatin loci in YFP-LacI-MCM2 expressing control and Orc6-depleted cells. YFP-LacI as a negative control. A representative experiment (n = 2) is shown. Positive cdc45 recruitment in YFP-LacI, 5.3% (n = 19); in YFP-LacI-MCM2 control, 52.8% (n = 36); in YFP-LacI-MCM2 Orc6-si, 27.5% (n = 40). Scale bar, 15µm. (E) Representative images of DNA fiber (upper panel). Percentages of restart tracks in total tracks counted (lower panel). Mean ± SD, n = 3. *p < 0.05 by unpaired two-tailed Student’s t test. See also Figure S3.

Next, we directly tested the function of hOrc6 in DNA replication by using DNA combing assay. Active replication forks were labeled by incorporation of 5-chloro-2’-deoxyuridine (CldU) followed by 5-iodo deoxyUridine (IdU), and the DNA fiber length was measured to determine fork movement. We did not observe any change in the fork velocity in control and hOrc6-depleted cells (Figure S3A). These results together imply that hOrc6 is required for Cdc45 association to MCM, and hence facilitates efficient helicase activation and S phase entry.

### Loss of Orc6 sensitizes cells to DNA damage and Orc6-depleted cells fail to activate ATR in response to replicative stress

Cells with replication defects often show reduced tolerance toward replication stress and DNA damage. Because we observed slower progression of cells through S phase upon Orc6 depletion (Figure 2I and 2J), we examined the replication fork dynamics in hOrc6-depleted cells upon replication stress treatment. Cells were first labeled with CldU for 30 min, treated with hydroxyurea for 2 hrs to induce stalling of replication fork, and subsequently released into the fresh medium containing IdU for another 30 min. DNA combing assay revealed that the cells lacking hOrc6 showed a significant and consistent decrease in fork restart (Figure 3E). This result implies that the slower S phase progression observed in hOrc6-depleted cells could be attributed to defects in the fork restart.

To test if the slowed progression through S phase resulted in replication stress and/or DNA damage, we performed comet assay to assess the extent of DNA breaks in hOrc6-depleted cells. There were no double- or single-strand breaks in the absence of hOrc6 under unperturbed condition. However, hOrc6-depleted cells in the presence of DNA damaging agents, including camptothecin (CPT) and hydrogen peroxide (H_2_O_2_) showed increased level of DNA damage compared to WT cells, as observed by both alkaline and neutral comet assays (Figure 4A and 4B). Furthermore, hOrc6-depleted or hOrc6-knockout cells were sensitive to DNA damage, as observed by significant nuclear fragmentation and decreased cell survival post-H_2_O_2_ treatment (Figure 4C and 4D). We reasoned that in the absence of hOrc6, the cells either fail to repair the damage or fail to sense DNA damage.

**Figure 4.**
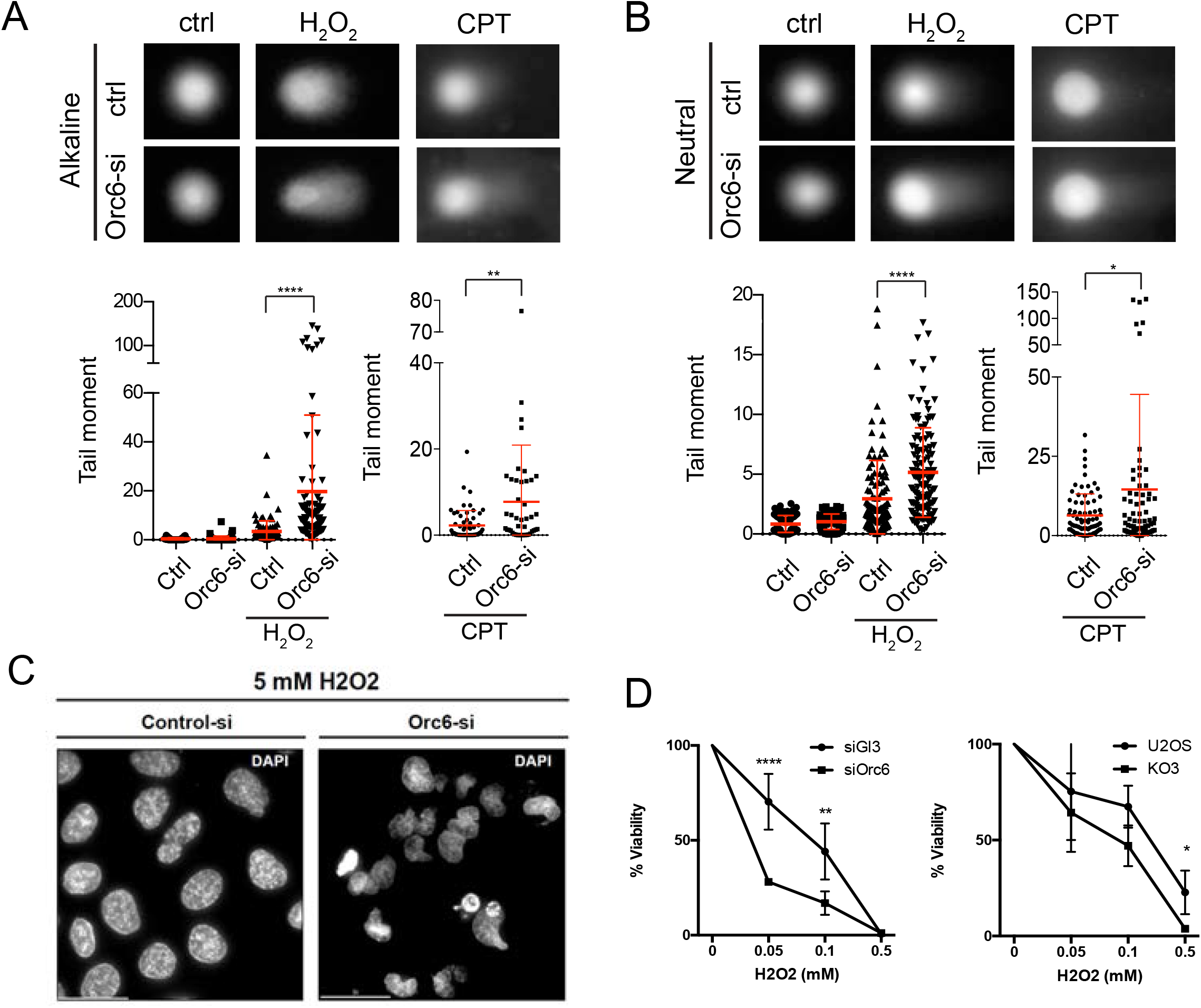
Depletion of Orc6 sensitizes cells to DNA damage. (A) Alkaline comet assay. Representative images are shown at top. One representative experiments for each is shown for H_2_O_2_-treated (n = 5) and CPT-treated (n = 2). Mean ± SD. **p < 0.01, and ****p < 0.0001 by unpaired two-tailed Student’s t test. (B) Neutral comet assay. Representative images are shown at top. Mean ± SD. *p < 0.05, and ****p < 0.0001 by unpaired two-tailed Student’s t test. (C) DAPI staining of control or Orc6 knockdown cells treated with H_2_O_2_. Scale bar, 30µm. (D) Clonogenic survival assay of H_2_O_2_-treated control and Orc6 knockdown cells (left); control and Orc6 knockout cells (right). Mean ± SD, n = 3. *p < 0.05, **p < 0.01, and ****p < 0.0001 by unpaired two-tailed Student’s t test.

We treated U2OS cells (with or without Orc6) with various DNA damaging agents, including H_2_O_2_ (oxidative damage), Neocarzinostatin (Radiomimetic drug), and HU for 4 hrs (to induce ssDNA and replication stress due to fork stalling) or for 24 hrs (to induce fork collapse and DSBs), and monitored the activation of ATR and ATM. The cells lacking Orc6 failed to activate ATR, as evident by decreased phospho-Chk1 and phospho-RPA, in response to different DNA damaging agents (Figures 5A, 5B and S4A). However, the ATM remained active as evident by intact Chk2 phosphorylation in both control and hOrc6-depleted cells. On the other hand, depletion of the other ORC subunits (Orc1 and Orc2) did not show significantly impaired ATR activation upon DNA damage (note* pRPA32) (Figure 5B). DNA-damaged cells lacking hOrc6 also showed reduced chromatin-association of RPA as observed by immunofluorescence staining using RPA antibodies following pre-extraction procedures (Figure 5C).

**Figure 5.**
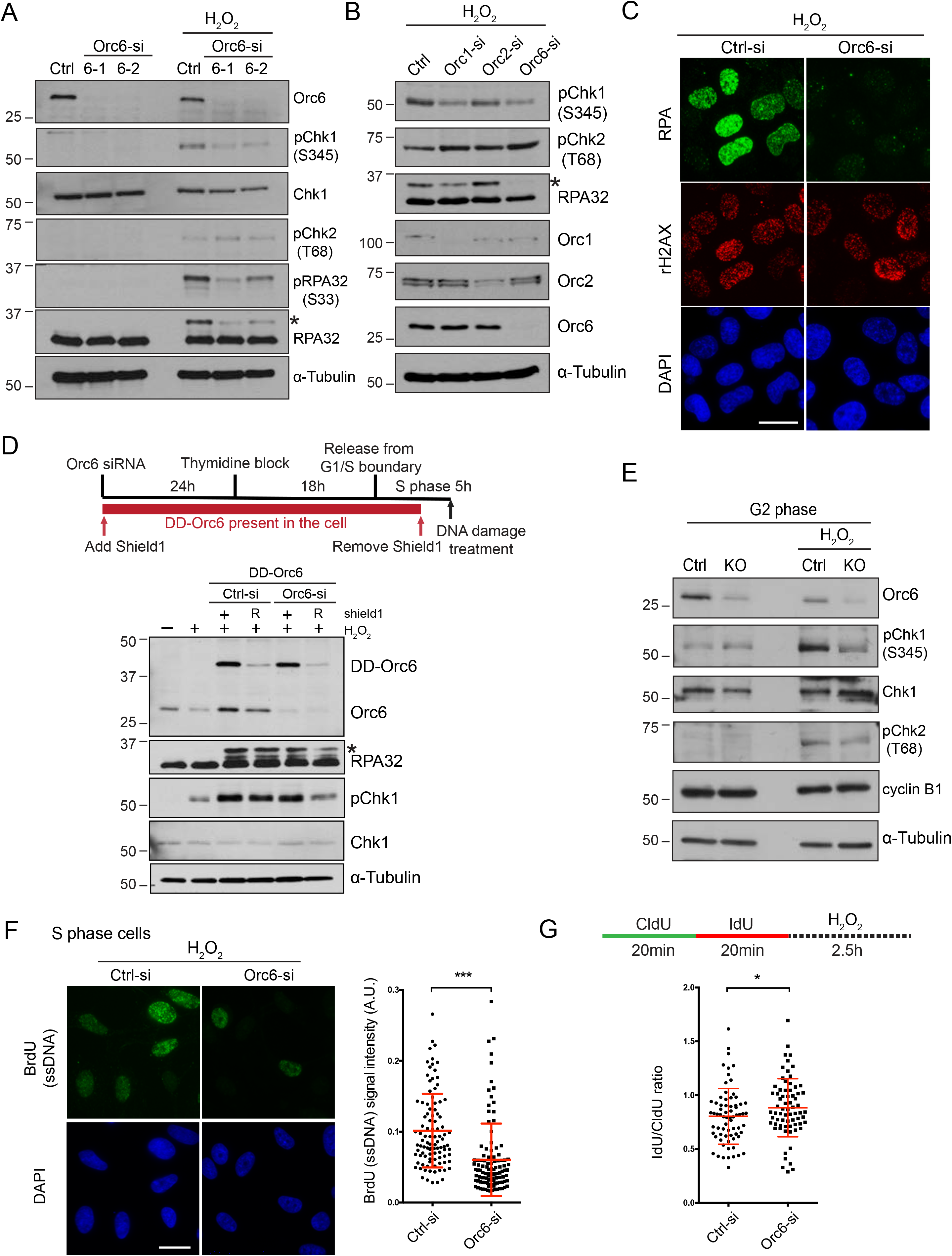
Loss of Orc6 leads to compromised ATR signaling pathway due to reduced ssDNA generation. (A) Western blot analysis of ATM/ATR pathway in control and Orc6 knockdown cells. *Asterisk* indicates hyperphosphorylation of RPA32. (B) Western blot for Orc1, Orc2 and Orc6 knockdown cells. (C) Immunostaining of chromatin-associated RPA and rH2AX in control and Orc6-depleted cells treated with H_2_O_2_. Scale bar, 25µm. (D) Upper panel: Schematic of the protocol for specifically depleting Orc6 in S phase and collecting DNA damage samples. Shield1 was removed 1h before H_2_O_2_ treatment and cells were collected 1h after treatment. Lower panel: western blot for analyzing ATR activation. R, removal of Shield1. (E) Western blot of G2 synchronized control and Orc6 knockout cells. Cyclin B1 as G2 marker. (F) Native BrdU staining for visualizing ssDNA (right) and quantification results (left). Scale bar, 25µm. A representative experiment (n=3) is shown. Mean ± SD; n > 100 for each group. ***p < 0.001 by unpaired two-tailed Student’s t test. (G) DNA fiber assay to determine nascent DNA resection/degradation after fork stalling. Higher IdU/CldU ratio indicates less resection, hence less ssDNA generation. A representative experiment (n = 2) is shown. Mean ± SD. ****p < 0.0001 by unpaired one-tailed Student’s t test. See also Figure S4, S5 and S6.

We next examined if the defects in RPA phosphorylation and Chk1 phosphorylation in cells lacking hOrc6 are a result of defective replication, or if they were due to a direct role of hOrc6 in ATR pathway. We synchronized the DD-Orc6-expressing cells (control as well as endogenous Orc6-depleted) in S phase by thymidine block, and subsequently degraded DD-Orc6 by removing Shield1 from the medium after the cells were released into S phase. We then induced DNA damage in the control and hOrc6-depleted S phase-synchronized cells (Figure 5D) and determined the status of ATR activity. We continued to observe defective ATR activation, as evident by reduced pChk1 and pRPA in Orc6-depleted cells that had been accumulated in S phase (Figure 5D). In addition, we synchronized control and hOrc6 knockout cells (hypomorph) into G2 phase and treated them with DNA damaging agents. The control G2 cells showed robust Chk1 and RPA phosphorylation whereas the Orc6-KO G2 cells failed to activate ATR, suggesting that hOrc6 facilitates ATR activation and that ATR activation defects in Orc6-depleted cells are not because of cell cycle effects (Figure 5E). These results support our model that hOrc6 is required for ATR activation and that the defects observed in Orc6-depleted cells are not simply a reflection of cell cycle defects.

The defect in RPA32 phosphorylation in hOrc6-depleted cells indicates that Orc6 may be involved in the upstream steps of the ATR signaling pathway. Processing of different DNA damages by various repair pathways yields ssDNA as a critical repair intermediate, which also serves to activate ATR. The two following possibilities could attribute to defects in ATR signaling upon hOrc6 depletion. First, hOrc6 could function in the recruitment of ATR signaling proteins to RPA-ssDNA platform. Secondly, hOrc6 could play a role in generating ssDNA after DNA damage. To test if hOrc6 plays a direct role in ATR activation, we performed an *in vitro* assay to examine the recruitment of ATR signaling proteins onto RPA-ssDNA platform in the presence or absence of hOrc6. We pre-assembled RPA-ssDNA complex by mixing recombinant RPA with 3’-biotinylated 70-mer single-strand DNA. We captured the complex using streptavidin-coated magnetic beads. These RPA-ssDNA structures were then incubated with nuclear extracts from control and Orc6-depleted U2OS cells, and we retrieved the proteins that bound to the structure. Western blotting was used to determine the effect of ATR signaling proteins (RPA32, pRPA32, ATRIP, Chk1, pChk1) recruitment and activation with or without hOrc6 (Figure S4B). We did not observe any defect in the recruitment of all of the proteins in Orc6-depleted cells. We also determined if hOrc6 plays any role in the RPA association to ssDNA. We performed ssDNA *in vitro* pull down with hOrc6 first being added to ssDNA then RPA, or vice-versa (Figure S4C). Again, these experiments demonstrated that hOrc6 did not have a direct role in facilitating RPA association to ssDNA. Based on these experiments, we concluded that hOrc6 does not play any direct role in the recruitment of signaling proteins to ssDNA and subsequent ATR activation.

Next, we evaluated whether hOrc6-depleted cells compromised the levels of ssDNA in cells. Towards this we quantified the levels of ssDNA in control and hOrc6-depleted cells using BrdU staining under non-denaturing conditions. DD-Orc6 cell line was used to ensure that both control and Orc6 knockdown cells were synchronized in S phase and had incorporated equal amount of BrdU. We observed a significant reduction in ssDNA in cells lacking Orc6 (Figure 5F). Based on these results, we conclude that hOrc6 is required for the generation of ssDNA post-DNA damage.

Our results suggest that the inability to generate ssDNA in hOrc6-depleted cells contributes to the defects in ATR activation. Many repair pathways require the generation of ssDNA, such as end resection after DSBs to allow homologous recombination. During excision repair and mismatch repair pathways, ssDNA is also generated in a regulated manner by the removal of damaged bases/strands (Yan et al., 2014). Since we observed hOrc6 at the replication fork, we set out to determine the efficiency of ssDNA generation at the fork in cells lacking Orc6 under DNA damage condition. Cells labeled sequentially with CldU and IdU (20 mins each) were immediately treated with DNA damaging agent. The resection/degradation of nascent DNA strand post-DNA damage was then calculated by measuring the length of the IdU-labelled fiber in DNA fiber assay. We observed that DNA-damaged cells lacking hOrc6 displayed significantly longer IdU tracks, indicating defects in nascent DNA degradation (Figure 5G). This result further confirms that the inability to generate ssDNA in hOrc6 depleted cells causes defects in ATR activation.

### Orc6 is upregulated and phosphorylated in response to oxidative stress

In yeast, Orc6 is phosphorylated in a cell cycle-dependent manner, with increased Orc6 phosphorylation as the cells exit G1. The hyperphosphorylation of yOrc6 following START is one of the known mechanisms by which cells prevent re-replication (Nguyen et al., 2001). We have observed that a population of hOrc6 is phosphorylated during DNA damage. We successfully mapped the DNA damage-mediated phosphorylation site of human Orc6 at T229 by performing PhosTag gel electrophoresis (Chakraborty et al., 2014) (Figure S5A).

By generating an antibody that specifically recognized only the pT229-modified version of hOrc6, we further investigated this particular phosphorylation. Orc6-pT229 occurred prominently when the cells were treated with oxidative or alkylating stress inducing agents such as okadaic acid, H_2_O_2_ and methyl methanesulfonate (MMS) (Figure S5B). Interestingly, cells showed enhanced Orc6-pT229 in response to oxidative stress, preferentially during S phase (Figure S5C). Both the total and the pT229-modified levels of Orc6 were elevated upon oxidative stress (Figures S5D-F). Treatment of cells with phosphatase (CIP) showed a loss of signal, confirming the specificity of the phospho-antibody (Figure S5D and S5E). Strikingly, Orc6-pT229 induced within 20 minutes of H_2_O_2_ treatment and persisted for ∼30 mins post-recovery from H_2_O_2_ release, following which the levels of Orc6-pT229 declined (Figure S5F). Meanwhile, the cells recovered from H_2_O_2_ continued to show enhanced levels of total Orc6.

We had previously demonstrated that cells lacking hOrc6 failed to activate ATR upon oxidative stress. Interestingly, the T229 of hOrc6 is an ATM/ATR consensus (TQ/SQ) site, implying that DNA oxidative stress-induced phosphorylation of Orc6-T229 could be mediated by ATM/ATR axis. In order to test whether the oxidative stress-induced Orc6-pT229 contributes to ATR activity, we generated HA-Orc6 U2OS cell lines stably expressing phospho-dead (T229A) or phospho-mimic (T229E). We found that ATR activity (pChk1 & pRPA32) post-oxidative stress was partially rescued by both wild type and T229E mutant of Orc6-expressing cells but not the cells that expressed the T229A mutant, further supporting the importance of pT229 of Orc6 in DDR (Figure S6A). It is critical to note that T229 is located adjacent to the ‘YxxWK’ conserved motif within the C-terminus of hOrc6, mutation of which is reported to impede the recruitment of Orc6 to ORC, and is linked to the Meier-Gorlin syndrome (Bleichert et al., 2013). We therefore tested whether the hOrc6-pT229 affected Orc6’s DNA binding activity and its interaction with other ORC subunits. We observed that both T229A and T229E mutants bound to DNA more efficiently than even the WT-Orc6 (Figure S6B) as observed by SiMPull assays. Also, the WT and both the mutants showed comparable levels of interaction with Orc2 (Figure S6C). These results imply that hOrc6-pT229 does not contribute to the DNA binding activity of Orc6, and also its ability to interact with ORC.

### Orc6 associates with the MMR complex by interacting with MutSα

To gain mechanistic insights into the role of hOrc6 in S phase progression and in DNA damage response, we identified hOrc6-interacting proteins during S phase (in presence or absence of DNA damage) by performing immunoprecipitation followed by mass spectrometry. The hOrc6 interacting-proteome revealed known interactors as well as other previously unreported interactions (Table S1). Strikingly, we found that in S phase extract, hOrc6 interacted with all the components of the MutSα (MSH2 and MSH6) and MutLα (MLH1 and PMS2) complexes that are known to initiate DNA mismatch repair (MMR) (Figure 6A). More importantly, H_2_O_2_-treated cells revealed enhanced interaction between hOrc6 and members of the MMR complex (Figure 6A). We validated the *in vivo* association between hOrc6 and MSH2 using PLA. The interaction between hOrc6 and MSH2 was enhanced in oxidative stress-induced cells (Figure 6B). Furthermore, the association between these two proteins became more significant in S phase cells (Figure 6B). To further examine the physical interaction of Orc6 with MutSα and MutLα, we conducted GST pull-down using purified proteins. We found that Orc6 directly interacts with MutSα but not MutLα (Figure 6C). Moreover, GST pull down using Orc6 truncations identified that Orc6 interacts with MutSα through its middle TFIIB (TFIIB-B) domain (Figure 6D and 6E). Our results indicate that hORC6 physically interacts with MutSα complex during S phase, and also during oxidative DNA damage.

**Figure 6.**
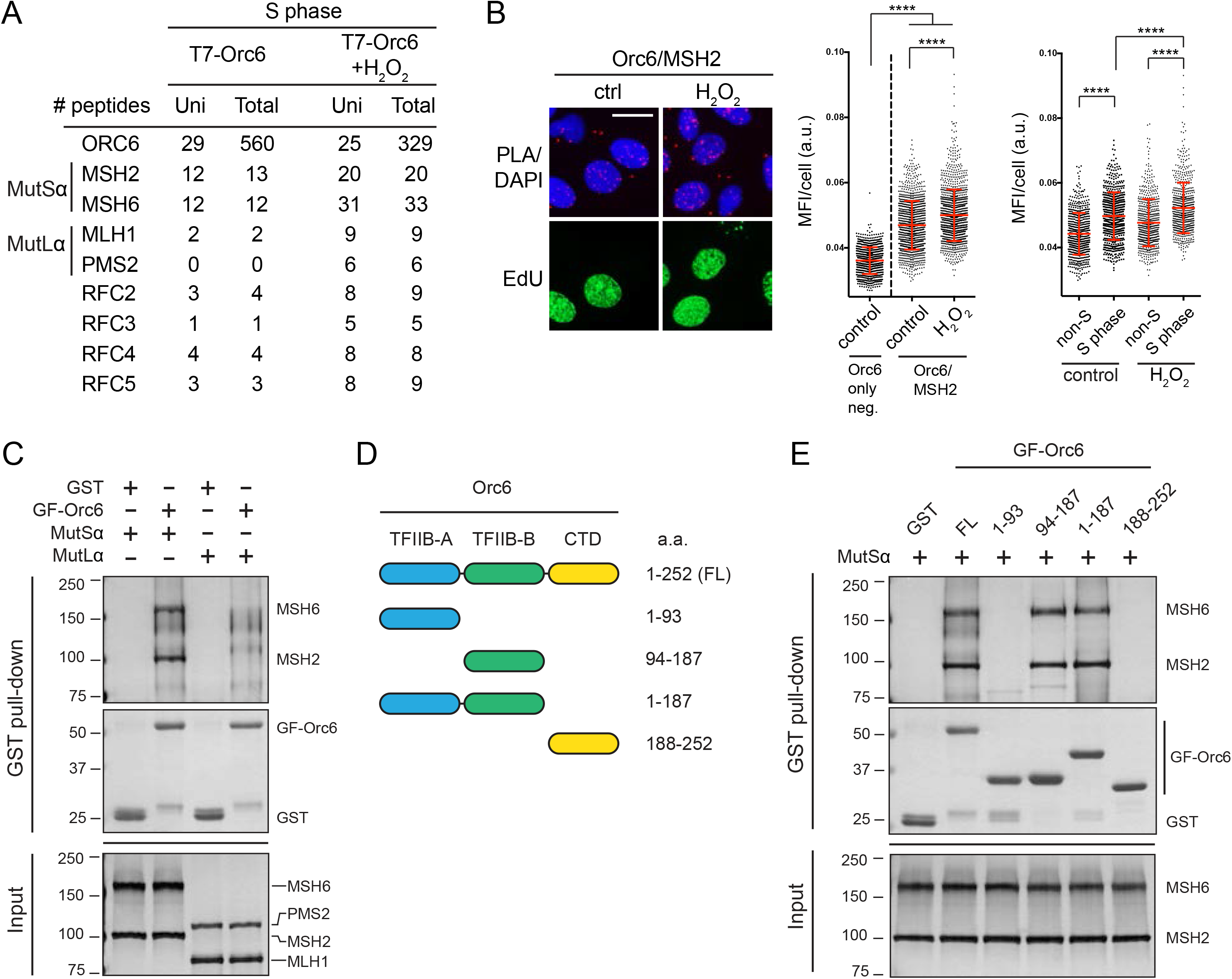
Orc6 interacts with MutSα using its middle TFIIB domain. (A) MMR proteins identified by IP-MS analysis from S phase synchronized U2OS cells expressing T7-Orc6, with or without H_2_O_2_ treatment. For each protein, numbers of unique peptides (Uni) and total peptides (Total) are presented. (B) Left panel: Orc6 and MSH2 association by PLA. EdU incorporation for determining S phase cells. Scale bar, 25µm. Middle panel: quantification of PLA. First column represents a negative control where MSH2 antibody was omitted. Right panel: further analysis of the quantification where S phase (EdU positive) and non-S phase (EdU negative) cells were separated in both control and H_2_O_2_ groups. Mean ± SD. ****p < 0.0001 by unpaired two-tailed Student’s t test. MFI: mean fluorescence intensity. a.u.: arbitrary unit. (C) Direct interaction of Orc6 with MutSα or MutLα examined by GST pull-down assay. Proteins on SDS-PAGE gels were visualized by silver stain (upper image and input) or Coomassie stain (middle image). GF-Orc6 stands for GST-Flag-Orc6. GST as a negative control. (D) Schematic illustration of Orc6 domains and different truncations. (E) Interaction of different truncations of GST-Flag-Orc6 with MutSα by GST pull-down assay. Proteins were visualized by silver stain (upper image and input) or Coomassie stain (middle image).

### Orc6 promotes MutLα recruitment to MutSα and facilitates MMR activity

The MMR process initiates when MutSα recognizes a mismatch on the daughter strand and recruits MutLα (Reyes et al., 2015). To determine the role of hOrc6 in the MMR process, we addressed the association of MutSα and MutLα in cells lacking hOrc6. Using PLA approach, we found that the interaction between members of the MutSα and MutLα was severely compromised (MSH6/MLH1 or MSH6/PMS2) in Orc6-depleted cells (Figure 7A). However, the interaction between components of MutSα complex itself (MSH2/MSH6) remained unaltered in cells lacking Orc6 (Figure 7B). Next, we determined the status of the chromatin association of individual members of the MutSα and MutLα complex in control and Orc6-depleted cells. DD-Orc6 cells were collected following the same protocol as in figure 5D to ensure cells are in S phase. Components of MutSα (MSH2 and MSH6) loaded onto the chromatin equally efficiently in control and Orc6-depleted (untreated as well as H_2_O_2_-treated) cells (Figure 7C). However, components of MutLα (MLH1 and PMS2) showed reduced chromatin association in H_2_O_2_-treated cells lacking Orc6 (Figure 7C). These data demonstrate that hOrc6 is required for efficient MMR complex assembly on chromatin during oxidative DNA damage.

**Figure 7.**
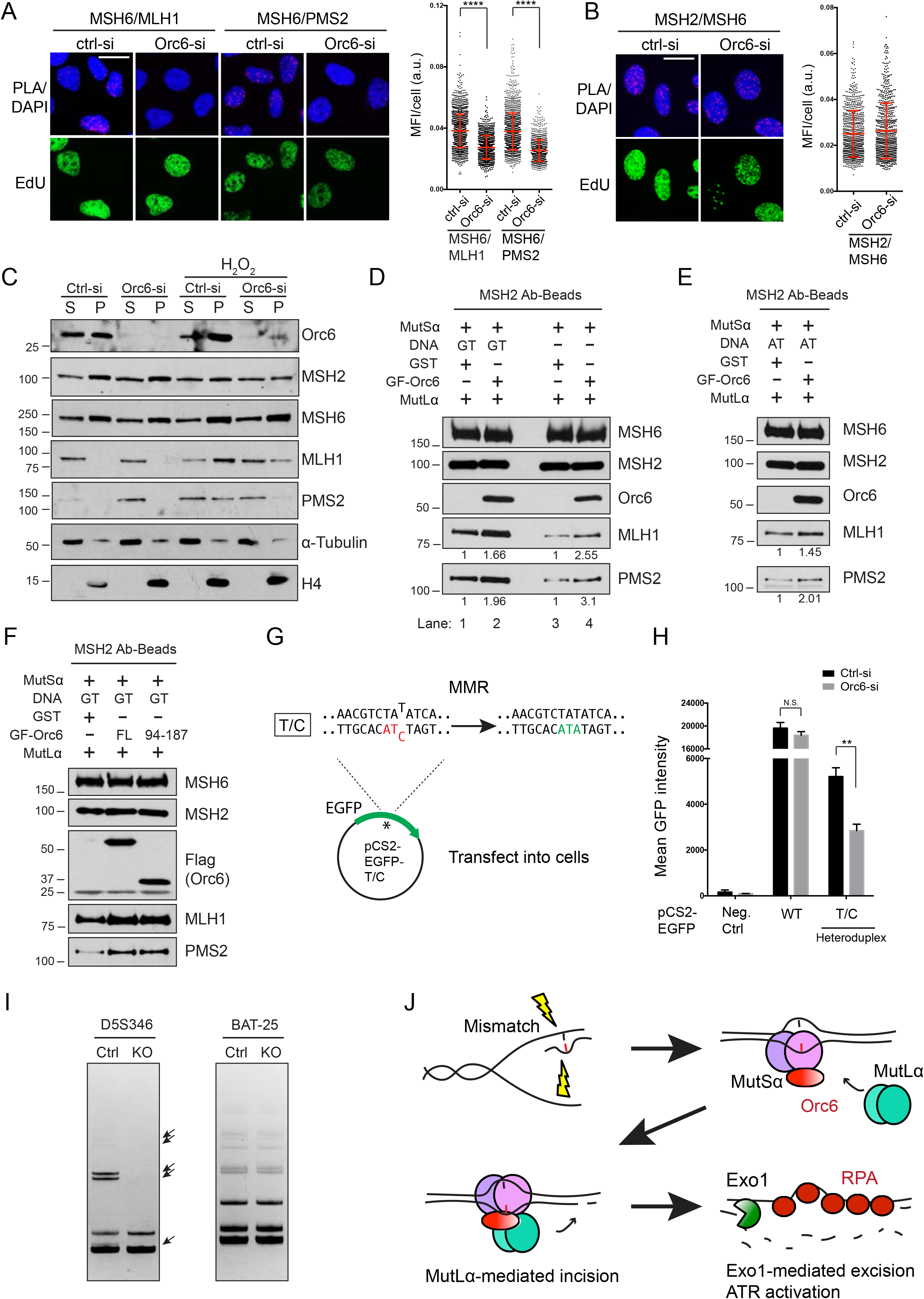
Orc6 facilitates MutLα recruitment to MutSα and promotes MMR efficiency. (A) Interaction between MSH6/MLH1 and MSH6/PMS2 quantified by PLA in control and Orc6-depleted cells. Representative images (left) and quantification (right) are shown. Scale bar, 25µm. Mean ± SD. ****p < 0.0001 by unpaired two-tailed Student’s t test. (B) Formation of MSH2/MSH6 complex (MutSα heterodimer) examined by PLA; representative images (left) and quantification (right) are shown. Scale bar, 25µm. Mean ± SD. (C) Western blot analysis of chromatin fractionation samples. The samples were prepared following the same protocol as in figure 5D. S, soluble; P, chromatin fraction. (D) Effect of Orc6 on the *in vitro* recruitment of MutLα to MutSα in the presence of G/T mismatch DNA (lane 1 and 2) or no DNA (lane 3 and 4). GF-Orc6 stands for GST-Flag-Orc6. GST as a control. (E) Effect of Orc6 on the *in vitro* recruitment of MutLα to MutSα in the presence of A/T homoduplex DNA. (F) *In vitro* recruitment of MutLα to MutSα using Orc6 full-length (FL) or TFIIB-B domain (94-187). (G) Illustration of the heteroduplex used in MMR assay. (H) Quantification of MMR activity. Mean GFP intensity was measured using flow cytometry. Mean ± SD, n = 3. **p < 0.01 by unpaired two-tailed Student’s t test. (I) Microsatellite loci D5S346 and BAT-25 were PCR-amplified from genomic DNA of wild-type U2OS control or Orc6 knockout (hypomorph) cells. The products were resolved on 15% polyacrylamide gels and stained with EtBr. Arrows indicate the differences observed. (J) Model for Orc6 function in facilitating the assembly of MMR complex for efficient repair and ATR activation. See also Figure S7.

To further address the mechanism of how hOrc6 functions in MMR complex assembly, we tested if the Orc6 promotes the association between MutLα and MutSα. We performed in vitro co-immunoprecipitation assay to determine the level of MutLα recruitment to immobilized MutSα using purified proteins in presence or absence of DNA. We observed that in the presence of Orc6, MutLα binds more efficiently to MutSα (Figure 7D and 7E). It is known that the affinity between MutSα and MutLα strongly increases in the presence of mismatch-containing DNA, which we have observed in our co-IP (Figure 7D, lane 1 and lane 3). However, the enhanced affinity between MutSα and MutLα by Orc6 is DNA and mismatch-independent, as we found the Orc6 facilitated the binding of MutLα to MutSα in all the experimental settings containing either heteroduplex DNA, no DNA or homoduplex DNA (Figure 7D and 7E). Moreover, the TFIIB-B domain of Orc6 is sufficient to promote MutLα binding to MutSα (Figure 7F). Therefore, these data suggest that Orc6 is an accessory factor which by binding to MutSα increases the affinity of MutLα to MutSα.

Having established that hOrc6 plays a role in MMR complex assembly and that in the absence of hOrc6, MutLα is recruited inefficiently, we set out to address if hOrc6 promotes MMR activity. To this end, we performed the *in vivo* MMR assay, whereby we determined the reversion of a mutated codon within EGFP and quantified the activity by measuring EGFP signal in the cells (Traver et al., 2015). We prepared a heteroduplex plasmid containing a mis-pair in the EGFP codon, where the sense strand is wild-type EGFP and the antisense strand has a mutation, which results in a premature stop codon (Figure 7G and S7A). Therefore, only when cells were able to repair the mismatch on the antisense strand could they express full-length wild-type EGFP and give fluorescence signal. Using this assay, we quantified the extent of MMR activity in WT U2OS cells that is MMR-proficient (Schopf et al., 2012) and compared it to Orc6-depleted U2OS cells. We observed that depletion of hOrc6 caused significant reduction in EGFP signal intensity in the T/C heteroduplex, suggesting that without Orc6, cells are less proficient in MMR (Figure 7H and S7B). Finally, to gain physiological view of hOrc6 function in the MMR, we investigated the microsatellite instability (MSI), a hallmark of defective MMR, in cells lacking hOrc6. For this purpose, Orc6 knockout U2OS cells (hypomorph) were sub-cultured to passage 16 (more than 30 divisions), and the genomic DNA was isolated for PCR analysis. By comparing Orc6 KO cells with wild-type U2OS, we observed alterations in the length of a dinucleotide-containing the microsatellite loci, D5S346, indicating that the loss of hOrc6 induces MSI, but not so for a mononucleotide-containing repeat (BAT-25) (Figure 7I). Our results collectively demonstrate that hOrc6 bound to MutSα facilitates the recruitment of MutLα to chromatin and thus is required for efficient MMR activity (Figure 7J).

## DISCUSSION

Primarily, based on studies from the yeast model system, Orc6, the smallest ORC subunit, is believed to function in DNA replication origin licensing and initiation. However, unlike other ORC subunits, function of human Orc6 is less clear due to its poorly conserved nature and conflicting biochemical data among species. In terms of replication licensing, recent structural studies have elucidated the requirement of yeast Orc6 in MCM loading (Li et al., 2018; Miller et al., 2019). However, it is still uncertain if human Orc6 functions the same way as in budding yeast. Our findings here suggest that hOrc6 is by and large dispensable for G1 origin licensing, indicating divergent roles for Orc6 in human and yeast licensing processes. Recently, using CRISPR approach to knockout human Orc1, Orc2, Orc5 and Orc2/Orc5, it was reported that human core ORC is dispensable for replication (Shibata and Dutta, 2020; Shibata et al., 2016). However, in human cells, acute depletion of Orc1 and Orc2 resulted in defects in the chromatin loading of MCMs. Nevertheless, in our experimental system, Orc6 behaved differently in the sense that its depletion did not alter MCM loading to the chromatin. Our observations of a significant S phase phenotype in the same experimental system strongly argue the important function of Orc6 after origin licensing.

Yeast Orc6 is also required after the licensing steps since depleting Orc6 after pre-RC formation has been shown to impair replication origin firing by destabilizing pre-RC (Semple et al., 2006). In human cells, we demonstrate that Orc6 associates with the replication fork and interacts with the components of the replication fork, the observation that are also supported by proteomic data (Alabert et al., 2014; Wessel et al., 2019). Using the DD-Orc6 to degrade Orc6 at specific stages of the cell cycle, we find hOrc6 to be dispensable for the licensing step, but essential for S phase progression. Our results argue that in human cells, Orc6 definitely functions after pre-RC formation. Most significantly, in cells lacking hOrc6, we find that Cdc45 loading onto chromatin was impaired. These results point to an inefficient CMG helicase activity and defective DNA replication progression in the absence of hOrc6. Without hOrc6, deficiency of Cdc45 chromatin loading results in fewer origins being activated, but the ones that are activated display normal fork progression rate.

The earliest steps of DNA replication, including the establishment of preRC are coordinated with the DNA damage network that prevents genomic instability (Cook, 2009). ATM (ataxia-telangiectasia mutated) and ATR (ATM and Rad3-related) are the key kinases that regulate DDR (DNA damage response) (Marechal and Zou, 2013). While ATM activation requires double strand DNA break bound by the MRN complex (Lee and Paull, 2005), ATR is typically activated when it senses single-stranded DNA coated by RPA (Marechal and Zou, 2013; Shiotani and Zou, 2009). Utilizing Stable Isotope Labeling by Amino acids in Cell culture (SILAC) quantitative proteomic approach, an earlier study identified over 700 ATM/ATR substrates upon IR-induced DNA damage, including hOrc6 at T229 site (Matsuoka et al., 2007). Multiple preRC factors have also been identified and studied. For example, Cdc6 physically interacts with ATR and this is thought to regulate the activation of replication checkpoint (Yoshida et al., 2010). Similarly, Mcm2 is an ATR substrate and Mcm2 associates with ATRIP and it is thought that the phosphorylation of MCM inhibits its DNA helicase activity or it may contribute to maintaining MCM at the stalled fork to prevent fork collapse (Cortez et al., 2004). Based on these data, it was proposed that a potential mechanism to block re-replication after DNA damage involves phosphorylation of preRC components by one or several of the checkpoint kinases. We observed that hOrc6 levels were elevated upon specific kinds of genotoxic stress like oxidizing agents, and it is phosphorylated in response to oxidative stress. We propose that the phosphorylated hOrc6 either acts as a brake or as a sensor of oxidative DNA damage to prevent replication progression. Future studies will test how phosphorylated hOrc6 inhibits replication progression in response to oxidative DNA damage.

Mismatch repair (MMR) is a process that recognizes and fixes errors during DNA replication. Our results indicate that hOrc6 is an accessary factor of the MMR complex, promoting MMR complex assembly and activity (Figure 7J). It is worth noting that although the structures of MutS and MutL are available, the interaction between them has been difficult to study (Friedhoff et al., 2016). During the eukaryotic MMR process, MutSα recognizes the mismatch and undergoes ATP-dependent conformational changes that allows the binding of MutLα. This MutSα-MutLα complex is very transient and dynamic. By using site-specific crosslinking, the transient *E. coli* MutS-MutL complex was successfully captured, providing valuable information about the MutS conformation when interacting with MutL (Groothuizen et al., 2015). However, the information about the human MutSα-MutLα structure is still lacking. It is also unclear if *in vivo* there is any additional factor influencing the MMR complex assembly. Our results that the association of Orc6 with MutSα increases the affinity of MutLα binding to MutSα is therefore of great importance for the MMR field. We propose that the binding of Orc6, as an accessory factor, stabilizes the MutSα at a conformation that allows MutLα to bind. Future structural studies are needed to investigate how hOrc6 influences MutSα structure.

During MMR, ssDNA is an important intermediate generated by EXO1 excision after the incision step (Constantin et al., 2005; Genschel and Modrich, 2003; Kadyrov et al., 2006; Zhang et al., 2005). More recently, it has been shown that MMR processing of a methylation-induced DNA lesion behind the replication fork causes ssDNA accumulation, interrupts fork progression, and might induce replication stress (Gupta et al., 2018). Thus, inefficient MMR activity could lead to reduced ssDNA generation. This is consistent with the defects in ssDNA generation that we observed in hOrc6-depleted cells. This is further corroborated by the fact that cells treated with oxidative or alkylating agents showed increased DNA damage without Orc6. Since there is not an efficient way to specifically induce mismatch on DNA, most studies utilize oxidative or alkylating agents. Base excision repair (BER) and nucleotide excision repair (NER) are believed to be the two main pathways that remove these lesions. However, studies suggest that MMR is also critical for the response to oxidative DNA damage (Bridge et al., 2014). Moreover, defects in MMR activity directly lead to reduced ATR signaling, as knockdown of MSH2 causes reduced Chk1 phosphorylation (Wang and Qin, 2003). Additionally, recognition of O6-meG/T mispairs by MutSα and MutLα directly recruits and activates ATR/ATRIP complex independent of RPA-ssDNA (Yoshioka et al., 2006). Thus, our finding that ATR is not fully activated in hOrc6-depleted cells is likely due to defective MMR repair pathway. On the other hand, *in vitro* studies have pointed that MMR is accomplished within 10-15 minutes (Constantin et al., 2005; Zhang et al., 2005), however *in vivo* it is likely to take longer depending on the amount and severity of the damage. We observe that phosphorylation of hOrc6 upon damage is rapid and the dephosphorylation is accomplished within 30 minutes, coinciding with the time needed for MMR. Together, our results point to a new regulatory mechanism in human cells whereby Orc6 travels along with replisome and acts as an accessory factor of MutSα upon encountering a mismatch, undergoes phosphorylation and promotes MMR complex formation to facilitate DNA damage repair and ATR activation.

It is well known that defects in MMR cause errors during DNA replication and are linked to a hereditary cancer syndrome, Lynch syndrome, often associated with microsatellite instability (Goellner, 2020). Further, colorectal tumors are often associated with defects in MMR (Li and Martin, 2016). It is worth noting that Orc6 levels are highly elevated in colorectal cancers (Xi et al., 2008), and the reduction of Orc6 sensitizes colorectal cancer cells to chemotherapeutic drugs (Gavin et al., 2008). Moreover, *ORC6* has been included as a predictor in three commonly used prognostic multigene expression profiles for breast cancer (Koleck and Conley, 2016). It has been known for a long time that the elevated level of Orc6 correlates with genome instability, yet the molecular details had remained to be elucidated. Our present studies provide novel insights into the role of Orc6 in the maintenance of genome integrity.

## Acknowledgements

We thank members of the Prasanth laboratory for discussions and suggestions. We thank Drs. M. Aladjem, J. Cook, A. Dutta, M. Mechali, B. Moriarity, S. Nair, B. Stillman, M. Wold, L. Zou for providing reagents and suggestions. We thank Dr. D. Rivier for critical reading of the manuscript. This work was supported by NSF-CMMB-IGERT fellowship to RH; NIH (R35 GM 122569) and NSF (PHY 1430124) awards to TH; NIH award (R01GM132128) to FAK; Cancer center at Illinois seed grant and Prairie Dragon Paddlers and NSF EAGER awards and NIH R01GM132458 and R21AG065748) to KVP; and NSF (1243372 and 1818286) and NIH (R01GM125196) awards to SGP. TH is an investigator with Howard Hughes Medical Institute.

## Author Contributions

Y-C.L. and A.C. designed, performed, and analyzed most experiments. D.L. performed MSI analysis; J.M. and R.H. did SiMPull experiments; L.Y.K. and F.A.K purified MutSα and MutLα proteins and prepared 5≠-nicked G-T and A-T DNAs; M.K.A. and S.A. helped with cloning, generating reagents and assistance with experiments. T.H. provided technical support and conceptual advice towards SiMPull experiments. F.A.K. provided reagents and technical suggestions towards biochemical co-IP and MSI analysis; S.G.P. and K.V.P. supervised the project. S.G.P. and Y-C.L. wrote the manuscript.

## Declaration of Interests

The authors declare no competing interests.

## STAR METHODS

### RESOURCES AVAILABILITY

#### Lead Contact

Further information and requests for resources and reagents should be directed to and will be fulfilled by the Lead Contact, Supriya Prasanth.

#### Materials Availability

All unique/stable reagents generated in this study are available from the Lead Contact with a completed Materials Transfer Agreement.

#### Data and Code Availability

The published article includes all datasets generated or analyzed during this study.

### EXPERIMENTAL MODEL AND SUBJECT DETAILS

#### Cell lines

Human cell lines U2OS and HEK293T were grown in DMEM containing high glucose and supplemented with 10% fetal bovine serum (FBS). Plasmids and siRNAs were delivered using Lipofectamine 2000 and Lipofectamine RNAiMax (Invitrogen), respectively. U2OS cells stably expressing HA-Orc6 or DD-Orc6 and Orc6 knockout cells were maintained in media containing selective antibiotics. Cell synchronization was done by nocodazole arrest for M and G1 phase samples, and by thymidine block for G1/S, S and G2 phase samples.

### METHOD DETAILS

#### Isolation of Proteins On Nascent DNA (iPOND)

A modified version of iPOND (Sirbu et al., 2013), described as aniPOND (Leung et al., 2013), was used. In brief, 1.5 x 10^8^ 293T cells were pulsed with 10µM of EdU for 10 min. For mature DNA samples, cells were chased with 10µM Thymidine for 1h. Cells were lysed in NEB (20mM HEPES pH7.9, 50mM NaCl, 3mM MgCl2, 300mM sucrose, and 0.5% NP-40) on ice for 15 min and nuclei were harvested by centrifugation. After washing with PBS, nuclei were incubated in freshly prepared click reaction cocktail (2 mM copper sulfate, 10 µM biotin-azide, and 100 mM sodium ascorbate in PBS) at 4 °C for 1h. Nuclei were then washed in PBS and resuspended in B1 buffer (25mM NaCl, 2mM EDTA, 1% NP-40 in 50mM Tris-HCl pH8). Next, sonication was performed at 4 °C using a bioruptor (Diagenode). Samples were then centrifuged at max speed for 10 min at 4 °C and supernatants were collected. An equal volume of B2 buffer (150mM NaCl, 2mM EDTA, 1% NP-40 in 50mM Tris-HCl pH8) was added to bring up the NaCl concentration, and input was taken at this point. EdU labeled DNA was then pulled down with 50µl of streptavidin beads (Dynabeads MyOne Streptavidin C1) at 4 °C overnight. Beads were washed with B2 buffer three times before boiled in laemmli sample buffer to elute captured proteins.

#### in Situ protein Interactions at Replication Forks (SIRF)

For SIRF experiments (Roy et al., 2018), cells were first pulsed with 125 µM EdU for 10 min. For Thymidine chases, cells were washed with PBS then 100 µM of Thymidine was added for 3h. Cells were then fixed in 2% paraformaldehyde (PFA) for 15 min at room temperature and permeabilized on ice with PBS containing 0.5% triton X-100. After washing with PBS, click reaction was performed with biotin-azide for 1h at room temperature. The coverslips were then blocked with blocking solution and preceded to standard PLA procedure using anti-biotin and antibodies indicated in the figures.

#### Proximity Ligation Assay (PLA)

PLA was performed using Sigma Duolink PLA as per the manufacture’s protocol. In brief, cells on coverslips were fixed in 2% PFA for 15 min at room temperature and permeabilized on ice with PBS containing 0.5% triton X-100. Coverslips were then blocked at 37 °C for 1h and incubated with primary antibodies overnight at 4 °C. After washing in buffer A, coverslips were incubated with Duolink PLA probes anti–mouse MINUS and anti–rabbit PLUS for 1h at 37 °C. Coverslips were then washed in buffer A and ligation was performed using ligation reaction mixture at 37 °C for 30 min. Next, coverslips were washed in buffer A and incubated in amplification reaction mixture for 100 min at 37 °C in the dark. Coverslips were subsequently washed twice in buffer B and once in 0.01x buffer B before DAPI staining.

For PLA experiments together with EdU labeling, 10 µM EdU was added 30 min before fixation. Click reaction using AF488-azide (Invitrogen) was performed after the permeabilization step and before the blocking step mentioned above, and all steps were performed in the dark.

#### Immunostaining

Cells were pre-extracted in 0.5% triton X-100 in CSK buffer (10mM PIPES pH 6.8, 100mM NaCl, 300mM sucrose, 3mM MgCl2) for 5 min on ice before fixation if needed. Cells were then fixed in 2% PFA for 15 min. If the pre-extraction was not performed, then cells were permeabilized on ice with PBS containing 0.5% triton X-100 for 5 min. Coverslips were then blocked in 1% normal goat serum (NGS) in PBS for 30 min and incubated with primary antibodies. Next, cells were washed in NGS/PBS and incubate in fluorophore-conjugated secondary antibodies for 45 min at room temperature. Cells were then washed in PBS and stained with DAPI.

For ssDNA visualization, cells were cultured in 10 µM BrdU for 36 h before any treatment to ensure same amount of BrdU incorporation. Cells were pre-extracted in CSK buffer containing 0.5% Triton X-100 for 5 min on ice, followed by fixing in 2% PFA for 20 min. Cells were washed with PBS and treated with chilled methanol for 15 min. Next, cells were treated with chilled acetone for 30 sec, washed with PBS again and blocked in PBST containing 2% BSA for 1 h. Cells were then incubated with the FITC-conjugated BrdU antibody at 4°C overnight. After washes with PBS, cells were stained with DAPI.

#### Immunoprecipitation

Cells were washed with PBS and lysed in IP lysis buffer (50mM Tris pH7.4, 150mM NaCl, 1mM MgCl2, 10% glycerol, 0.2% NP-40) containing protease inhibitors. Lysates were then sonicated and treated with benzonase nuclease (Sigma) for 30 min at room temperature, then EDTA was added to 2mM. Centrifugation was done at max speed for 10 min to remove insoluble debris. Next, lysates were pre-cleared with Gammabind G sepharose (GE healthcare Life Science) for 30 min at 4 °C. Antibodies were then added into lysates and incubated at 4 °C overnight. Proteins bound by antibodies were pulled down by Gammabind G Sepharose for 3h at 4°C. After incubation, beads were washed in lysis buffer and captured proteins were eluted and analyzed with western blot or mass spectrometry. Mass spectrometry and data analysis were performed by the Taplin Biological Mass Spectrometry Facility.

#### Flow cytometry

For PI cell cycle profile, cells were collected and washed once in ice cold PBS, resuspended in PBS + 1% NGS, and fixed in 90% chilled ethanol overnight. Cells were then washed and resuspended in PBS + 1% NGS with 120 µg/ml propidium iodide (PI) and 10 µg/ml RNase A for 45 min at 37 °C. DNA content was measured by flow cytometry. For BrdU-PI fow, cells were pulsed with BrdU for 30 min and stained with FITC-conjugated BrdU antibody before PI staining.

MCM-PI flow was done following methods previously described (Matson et al., 2017) with modifications. Briefly, cells were collected and pre-extracted with 0.5% triton X-100 in CSK buffer for 5 min on ice to remove soluble MCMs. Cells were then fixed in 1% PFA for 15 min RT, and stained with MCM3 antibody for 1h at 37 °C in 1% BSA/PBS with 0.1% NP-40. Next, Cells were incubated with fluorophore-conjugated secondary antibody for 1h at 37 °C. Finally, after washes, cells were stained with PI as indicated above.

#### Chromatin fractionation

U2OS cells were resuspended with solution A (10mM HEPES pH7.9, 10mM KCl, 1.5mM MgCl2, 0.34M sucrose, 1mM DTT, 10% glycerol and 0.1% Triton X-100) and incubate on ice for 5min. The cytoplasmic fraction (S2) was then separated from the nuclei by centrifuging at 4°C at 1400g for 4min. Isolated nuclei were then washed with solution A without Triton X-100. The nuclei pellet was resuspended with solution B (3mM EDTA, 0.2mM EGTA, and 1mM DTT) and incubated on ice for 30min. The nuclear soluble fraction (S3) was then separated by centrifuging at 4°C at 1700g for 4min. The S2 and S3 fractions can be combined as total soluble fraction (S). The isolated chromatin pellets were then washed with buffer B. Finally, the chromatin pellets were resuspended in solution A and sonicated for 1min to get P3 fraction.

#### Comet assay

Comet assay was performed using CometAssay Kit (Trevigen) following the manufacturer’s instructions. In brief, cells were collected by trypsinization, embedded in low-melting agarose and placed on CometSlides. After agarose solidifying, the slides were immersed in lysis solution for 30 min then subjected to electrophoresis for 30 min. For alkaline comet assay, the slides were incubated in alkaline unwinding solution before electrophoresis. After washing in water and 70% ethanol for 5 min each, the slides were allowed to dry and DNA was visualized using SYBR safe staining.

#### DNA fiber assay

Cells were labeled with 50mM CldU and 200mM IdU according to schemes in figures. DNA fibers were prepared on vinyl-silane coated coverslips using the FiberComb molecular combing system (Genomic Vision) as per the manufacture’s protocol. To visualize the CldU and IdU tracks, DNA fibers on coverslips were denatured in denaturation solution (0.5M NaOH, 1M NaCl) for 8min at room temperature. Coverslips were then washed with PBS and dehydrated in 70%, 90%, and 100% ethanol for 5min each. Coverslips were blocked with 1% BSA in PBST, followed by incubating in antibodies against CldU and IdU. After washing in BSA/PBST, the coverslips were incubated in FITC-conjugated goat anti-rat IgG and Texas Red-conjugated goat anti-mouse IgG. The images were captured using Zeiss Axiovision system.

#### Protein purification

Human RPA and MutSα were purified as described previously (Binz et al., 2006; Dzantiev et al., 2004). p11d-tRPA was a kind gift from Dr. Marc Wold (University of Iowa, Iowa City, IA). Human GST-Flag-Orc6 was induced for overexpression in E. coli BL21 (DE3) codon (+). The cells were collected and resuspended in GST buffer (50mM Tris pH7.5, 0.1mM EDTA, 150mM NaCl, 1mM DTT and 5% glycerol) containing lysozyme 0.5mg/ml and 0.1% Triton X-100 followed by sonication. The lysate was collected by centrifugation and the supernatant was subjected to GSTrap column (GE healthcare) using an AKTA pure system. After washing with GST buffer containing 500mM NaCl, GST-Orc6 was eluted in GST buffer containing 20mM reduced glutathione. Human Flag-MutLα was expressed in insect Sf9 cells and purified by chromatographies on α-Flag beads and MonoQ and MonoS columns.

#### GST pull down assay

GST control or GST-Flag-Orc6 was induced for overexpression in E. coli BL21 (DE3) codon (+), and lysate was prepared as described in protein purification part. The lysate was incubated with Glutathione-Agarose beads (Sigma) for 1h at 4°C. After protein binding, the beads were washed twice with GST buffer containing 500mM NaCl and once with GST buffer. For each GST pull down reaction, 2.5 µl of the packed beads containing 25 µg GST fusion protein was equilibrated in buffer A (20 mM HEPES pH 7.4, 100 mM NaCl, 0.1 mM EDTA, 0.01% Nonidet P-40, 5% glycerol, 0.1 mM DTT, and 0.2 mM PMSF). MutSα or MutLα were added as indicated in the figures to a final reaction volume of 30 µl containing buffer A. After incubated for 1h at 4°C, the beads were washed extensively with buffer A and bound proteins were eluted by boiling in laemmli sample buffer.

#### In vitro MutLα recruitment by co-immunoprecipitation

For each co-IP reaction, 5 µl packed GammaBind G sepharose beads were mixed with 5 µg of MSH2 antibody (Santa Cruz) overnight at 4°C. The beads with MSH2 antibody immobilized on were then equilibrated in MMR buffer (20 mM HEPES pH 7.4, 100 mM KCl, 5 mM MgCl2, 2 mM ATP, 4 mM DTT, 0.4 mg/ml BSA, 3–5% (v/v) glycerol), and MutSα was added in the final volume of 30 µl containing MMR buffer. The mixture was incubated for 2h at 4°C with gentle rotation. Afterward, the beads were washed, and 5≠-nicked G-T or A-T DNA (Dzantiev et al., 2004) and proteins (GST control/GST-Flag-Orc6 and MutLα) were subsequently added to the reaction mixture as indicated in the figures; the final reaction volume was 30 µl containing MMR buffer. After incubating for 1.5h at 4°C, the beads were extensively washed twice in MMR buffer containing 5% skim milk and once in MMR buffer. The proteins bound were finally eluted by boiling in laemmli sample buffer.

#### Mismatch repair (MMR) assay

In vivo mismatch repair efficiency is measured using MMR assay previously described (Traver et al., 2015). pCS2-EGFP-WT (wild type) was a kind gift from Dr. Marcel Méchali (Laboratory of DNA Replication and Genome Dynamics, Institute of Human Genetics, CNRS, Montpellier, France). pCS2-EGFP-t456g was generated from WT by Quikchange Site-Directed mutagenesis procedure. The pCS2-EGFP-WT plasmid was amplified into linear form with two primers, WT_Fwd and WT_Rev (see supplemental material), of which only the reverse one was 5’-phosphorylated. The mutated pCS2-EGFP-t456g was amplified with another set of primers, MIS_Fwd and MIS_Rev, of which only the forward one was 5’-phosphorylated. The PCR products were purified and digested using Lambda exonuclease (NEB) that degrades only the 5’-phosphorylated strands. The remaining single strands were then mixed (WT+t456g), denatured at 97°C for 5 min and annealed by slowly cooling to room temperature, allowing them to form nicked circular plasmids with T/C mismatch. The heteroduplex products were transfected into cells to observe mismatch repair efficiency by measuring EGFP signal.

#### Microsatellite Instability (MSI) Assay

U2OS Orc6 KO cells (clone 3) were cultured to passage 16 (more than 30 divisions) and then genomic DNA was extracted. Genomic DNA of wild-type U2OS cells was used as control. Genomic PCR was carried out under following conditions: initial denature at 98 °C for 5 min, followed by 35 cycles of 98 °C for 30 sec, annealing temperatures specific to each primer pair for 30 sec and 72 °C for 30 sec, with a final extension at 72 °C for another 7 min. PCR products were analyzed by 15% non-denaturing polyacrylamide gel electrophoresis and visualized by EtBr staining.

#### Single Molecule Pull down (SiMPull)

For replication fork-like DNA substrate preparation, partial duplexes of T1/P1 and T2/P2 were generated by annealing equimolar concentrations of the oligonucleotides in a buffer containing 10 mM Tris, pH 8.0 and 50 mM NaCl at 95 °C for 3 min, followed by slow cooling to room temperature. The two duplexes were then combined together and incubated overnight at room temperature to generate the replication fork probe. The reaction mixture was subjected to PAGE purification prior to use.

For double-strand DNA substrate, a T1-int/T2 duplex was generated similarly by annealing T1-int and T2 at 95 °C for 3 min, slow cooling to ambient temperature, followed by PAGE purification. The purified constructs were stored in buffer containing 10 mM Tris, pH 8.0 and 25 mM NaCl.

Single-molecule experiments were performed on a prism-type TIRF microscope equipped with an electron-multiplying CCD camera (EM-CCD) (Roy et al., 2008). For single-molecule pull-down experiments quartz slides and glass cover slips were passivated with 5000 MW methoxy poly- (ethylene glycol) (mPEG, Laysan Bio) doped with 2-5 % 5000 MW biotinylated PEG (Laysan Bio). Each passivated slide and cover slip was assembled into flow chambers. GST-Flag-Orc6 wild-type and its variants (T229A and T229E) were diluted to 1 nM and pulled down with biotinylated antibodies against GST (Abcam, ab87834), already immobilized on the surface via neutravidin-biotin linkage. Any unbound protein was washed off after 10 min incubation and a predefined concentration of DNA substrate (replication fork-DNA, dsDNA, i.e. T1-int/T2 or ssDNA, i.e. T1-int) was introduced in the imaging chamber. The protein-DNA complexes were imaged post 15 min incubation with the DNA substrate, in a buffer containing 20 mM HEPES (pH 8), 60 mM KCl, 5 mM MgCl2, 0.5 mM EDTA and 8% Glycerol. Cy3- and Cy5-tagged DNA were excited at 532 nm and 640 nm respectively and the emitted fluorescence signal was collected via band pass filters (HQ 570/40, Semrock for Cy3 and 665LP, Semrock for Cy5). 15 frames were recorded from each of 20 different imaging areas (5,000 µm2) and isolated single-molecule peaks were identified by fitting a Gaussian profile to the average intensity from the first ten frames. Mean spot-count per image for Cy3 and Cy5 was obtained by averaging 20 imaging areas using MATLAB scripts. All experiments were carried out at room-temperature.

For Orc6 interaction with RPA, SiMPull experiments were carried out in flow chambers prepared on quartz microscope slides, which were passivated with methoxy-polyethylene glycol (mPEG) doped with 1% biotin-PEG. Biotinylated HA antibody was immobilized on PEG passivated surfaces at approximately 20 nM concentration for 20 min after coating the flow chambers with 0.2 mg/ml NeutrAvidin for 5 min. Cells were collected 24 h after transient transfection, and lysed in high salt buffer (50 mM Tris-HCl pH 7.4, 500 mM NaCl, 10% glycerol, 0.25% Triton X-100 with protease inhibitors) at 4°C for 20 min. Equal amount of zero salt buffer were added and incubated for another 10 min. After centrifugation, the supernatants were used for SiMPull analysis. Samples were appropriately diluted with T50 buffer (10 mM Tris-HCl pH 8.0, 50 mM NaCl, 0.1 mg/ml BSA) to obtain optimal single molecule density on the surface. Diluted Samples were incubated in the chamber for 20 min and washed with the buffer.

#### Single-strand DNA in vitro pull down

To generate ssDNA for the in vitro binding assay, synthetic 70-mer DNA oligomers with 3’ biotinylation were attached to streptavidin-coated magnetic beads (Dynabeads™ MyOne™ Streptavidin C1) in binding buffer (10mM Tris pH7.5, 100mM NaCl, 10% glycerol, 0.01% NP-40, 10mg/ml BSA). Generally, 100 pmol of biotinylated DNA oligomers were incubated with 100 µl of dynabeads for 30 min at room temperature. The ssDNA-bound beads were washed with binding buffer to remove unbound DNA oligomers. For each reaction, 5 µl ssDNA-bound beads were used. For RPA coated ssDNA-beads, 25 pmol of purified RPA was incubated with every 5 µl of ssDNA-beads. The beads were used for in vitro pull down as indicated in the figures.

### QUANTIFICATION AND STATISTICAL ANALYSES

Microscopy image analyses and quantifications were done using CellProfiler (McQuin et al., 2018). Comet assay quantifications were done using OpenComet (Gyori et al., 2014). Statistical analyses were performed by two-tailed Student’s t test unless indicated differently in the figure legends. Quantifications represented as mean ± standard deviation. P value asterisk convention: * = p < 0.05, ** = p < 0.01, *** = p < 0.001, and **** = p < 0.0001. Further statistical details of experiments can be found in the figure legends.

### KEY RESOURCES TABLE

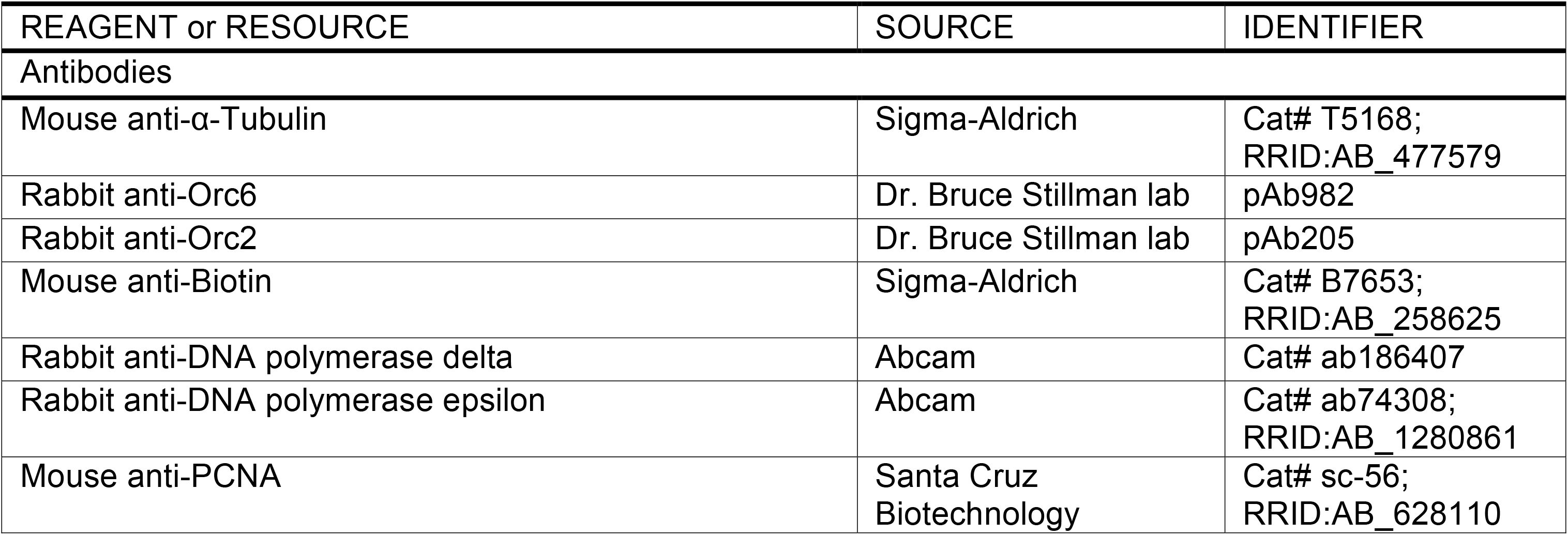
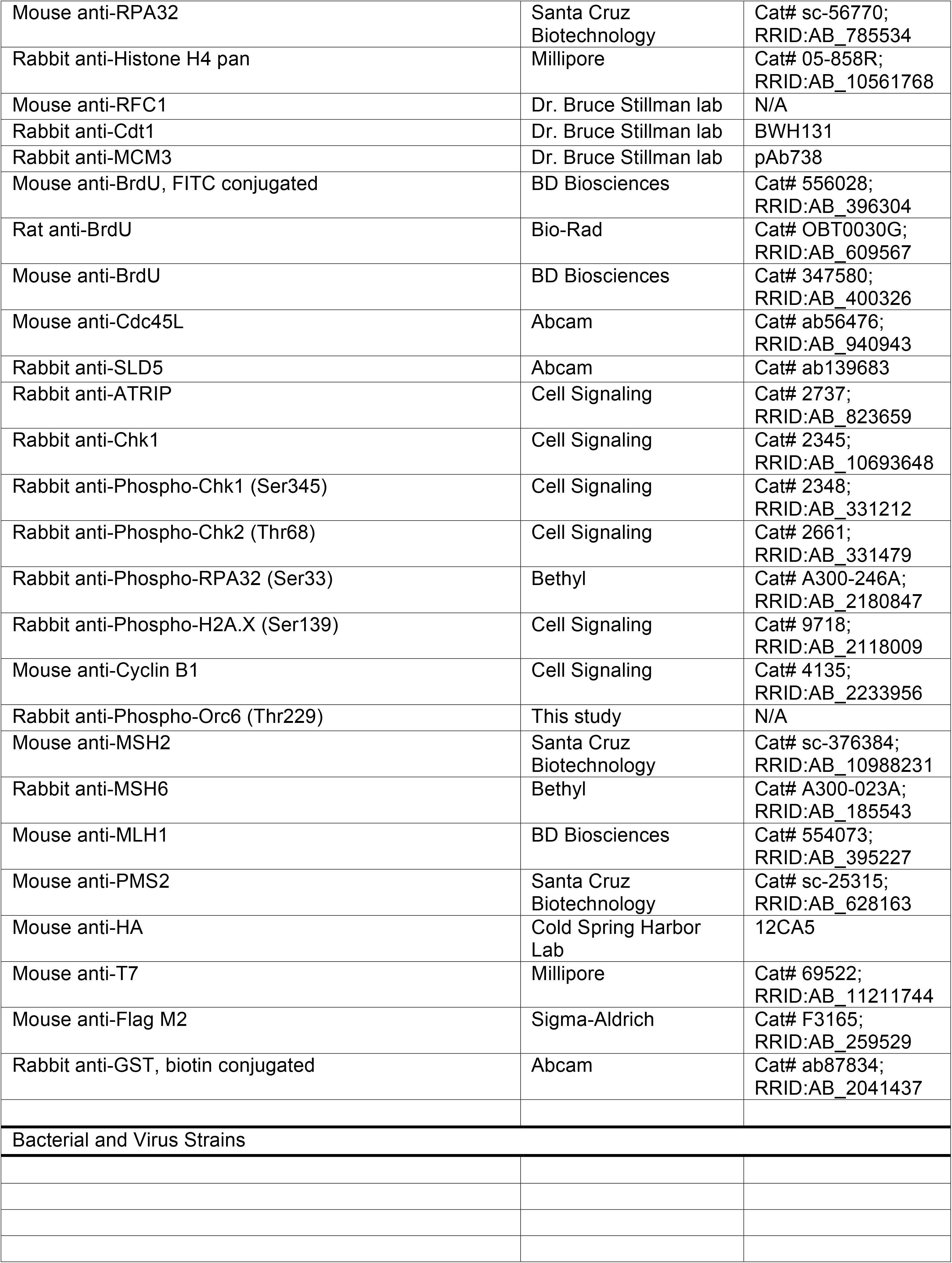
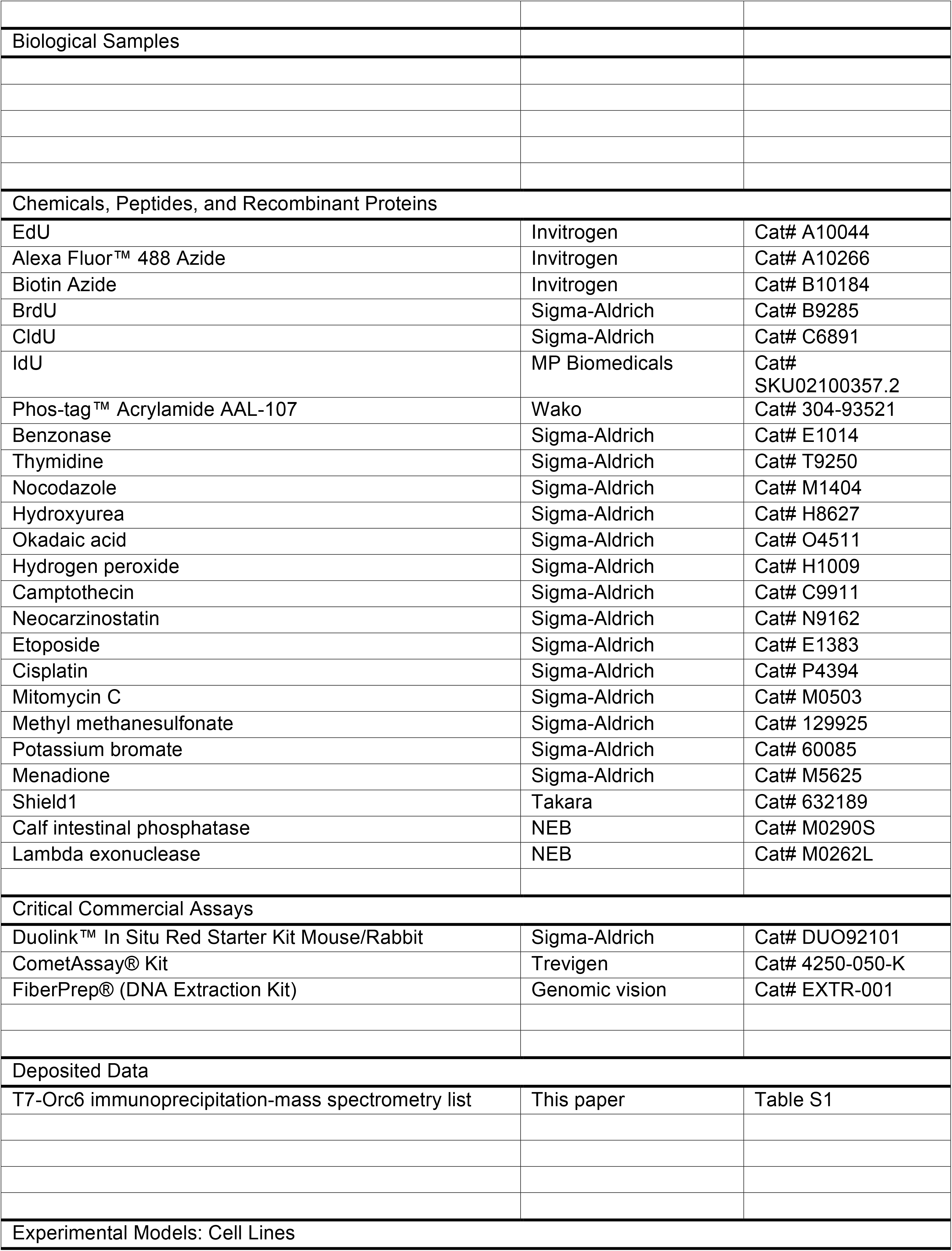
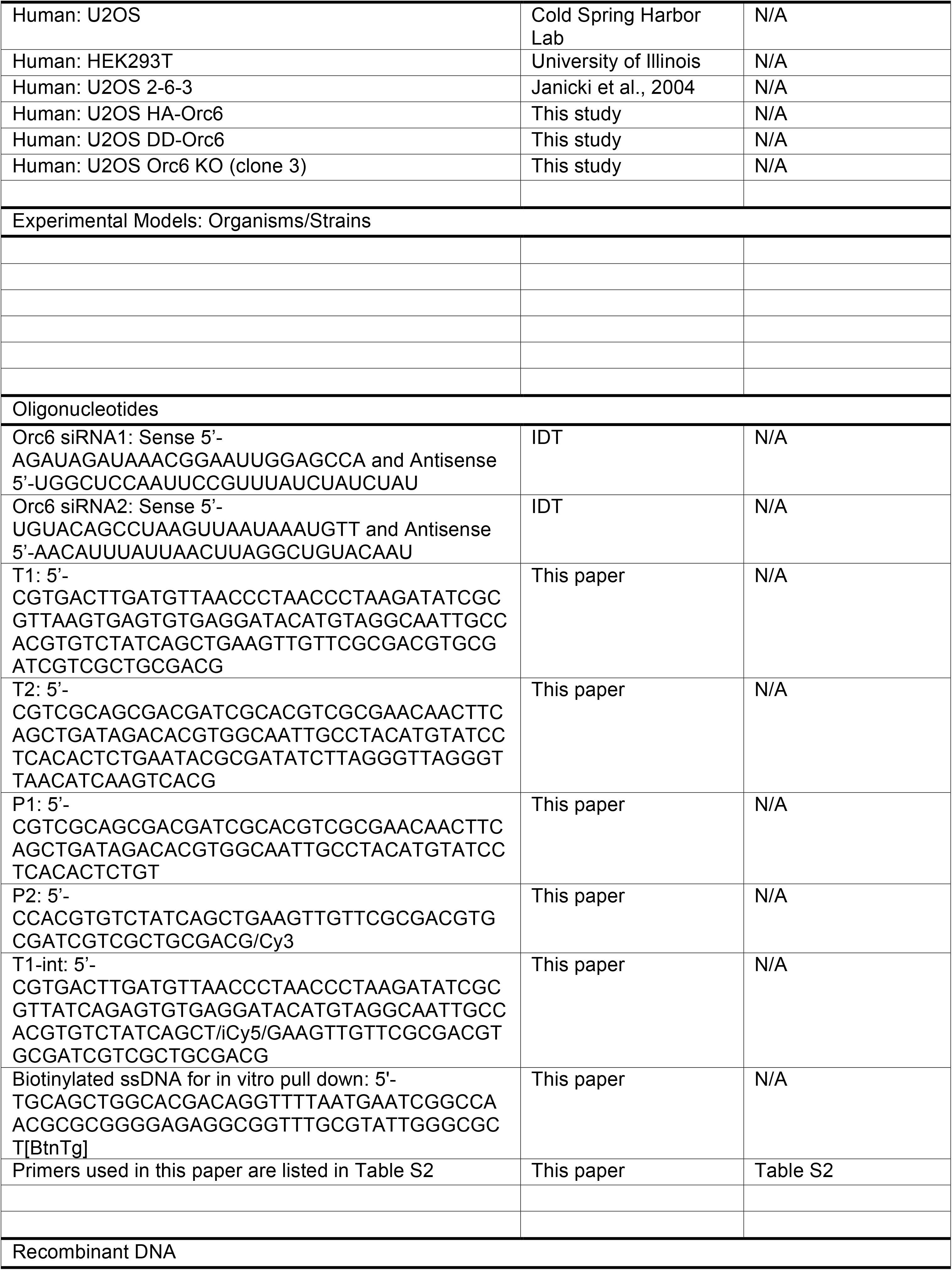
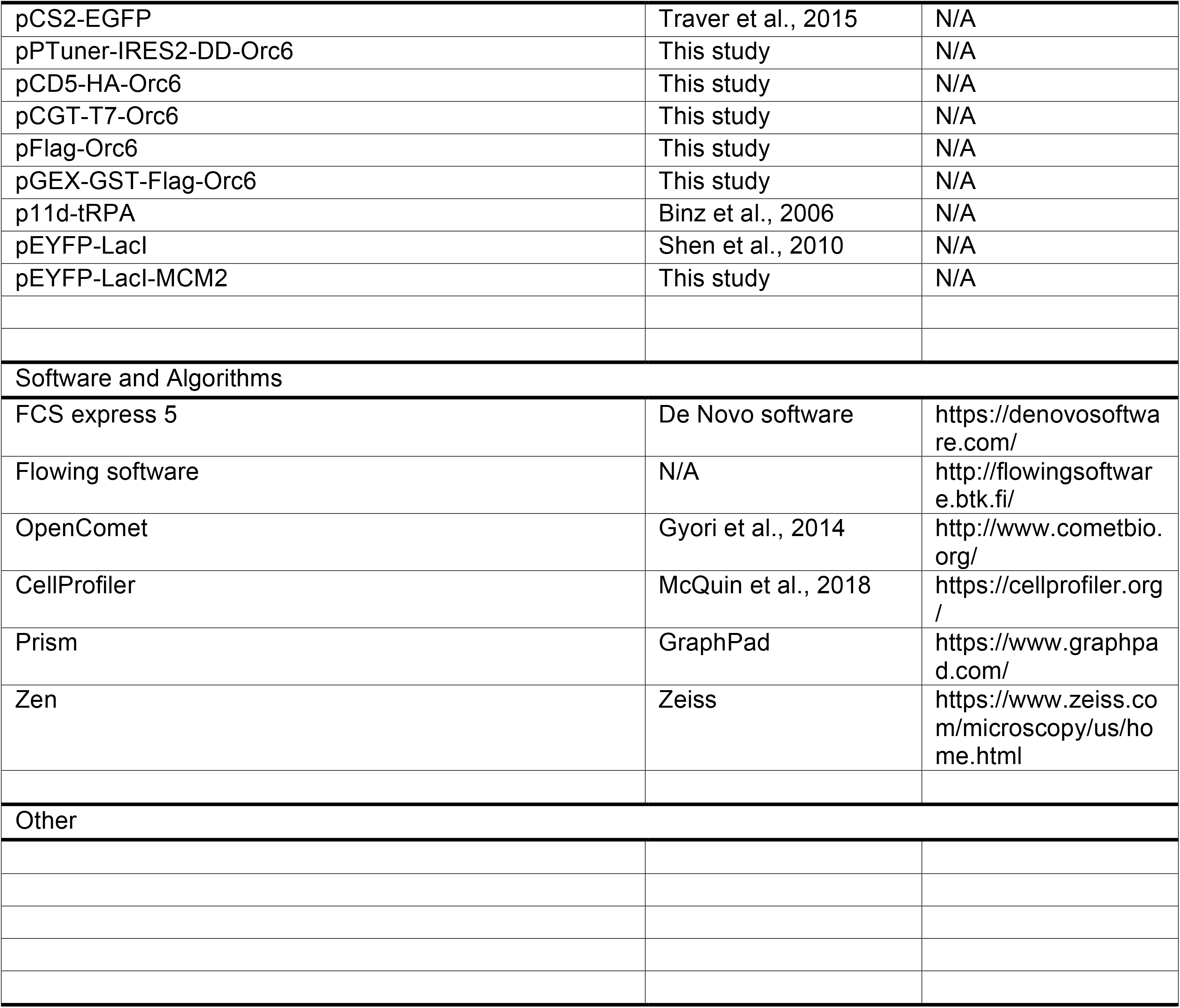

## SUPPLEMENTAL INFORMATION TITLES AND LEGENDS

**Figure S1.**
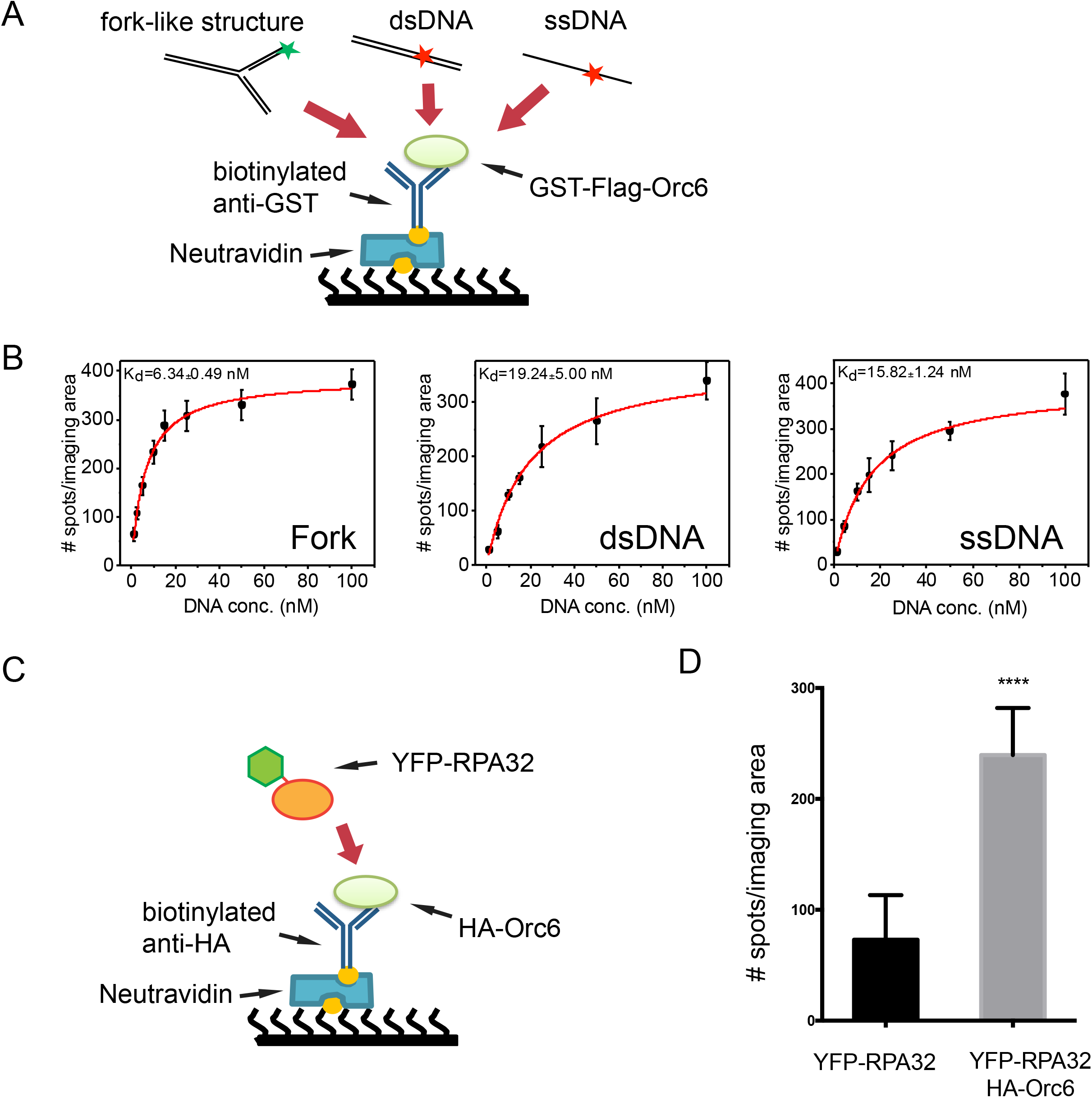
Orc6 preferably binds to fork like DNA structure and interacts with RPA (Related to figure 1) (A) Schematic illustration of SiMPull to test Orc6’s DNA binding ability to different DNA substrates. (B) Binding curves showing Orc6 binding affinity to different DNA substrates. (C) Schematic illustration of SiMPull to examine interaction between Orc6 and RPA32. (D) Quantification of Orc6 and RPA32 interaction. One signal spot represents one RPA molecule. Mean ± SD. ****p < 0.0001 by unpaired two-tailed Student’s t test.

**Figure S2.**
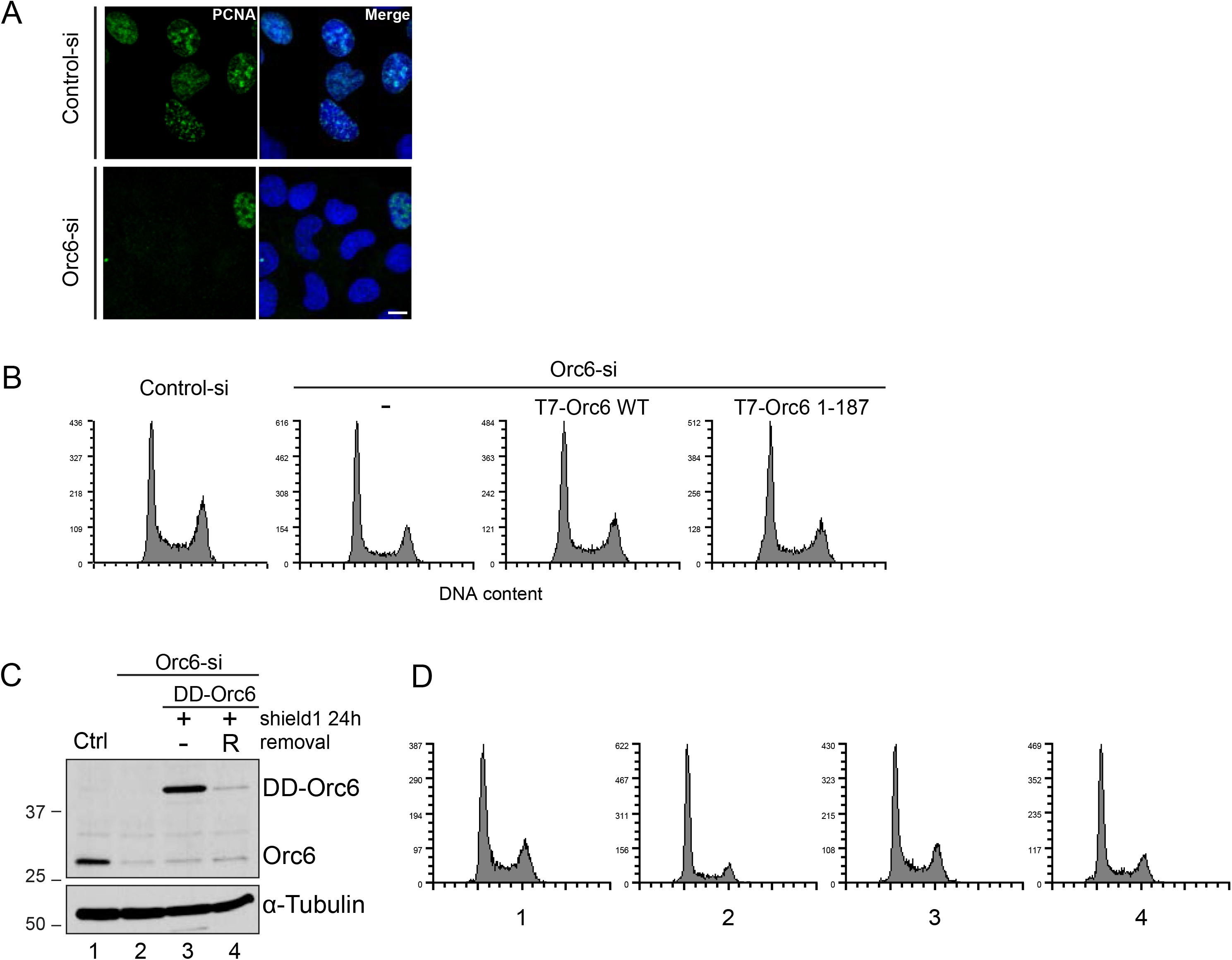
Orc6 knockdown and the validation of DD-Orc6 degron system (Related to figure 2) (A) Immunostaining analysis of Orc6 knockdown cells. PCNA staining was used to mark S phase cells. Scale bar, 15µm. (B) Cell cycle profile of endogenous Orc6-depleted cells substituted with tagged wild-type Orc6 or C-terminal truncated Orc6 (a.a.1-187). (C) Western blot showing the protein level of endogenous Orc6 and DD-Orc6 in the absence and presence of Shield1. (D) Cell cycle profile of samples from (C) by PI flow cytometry.

**Figure S3.**
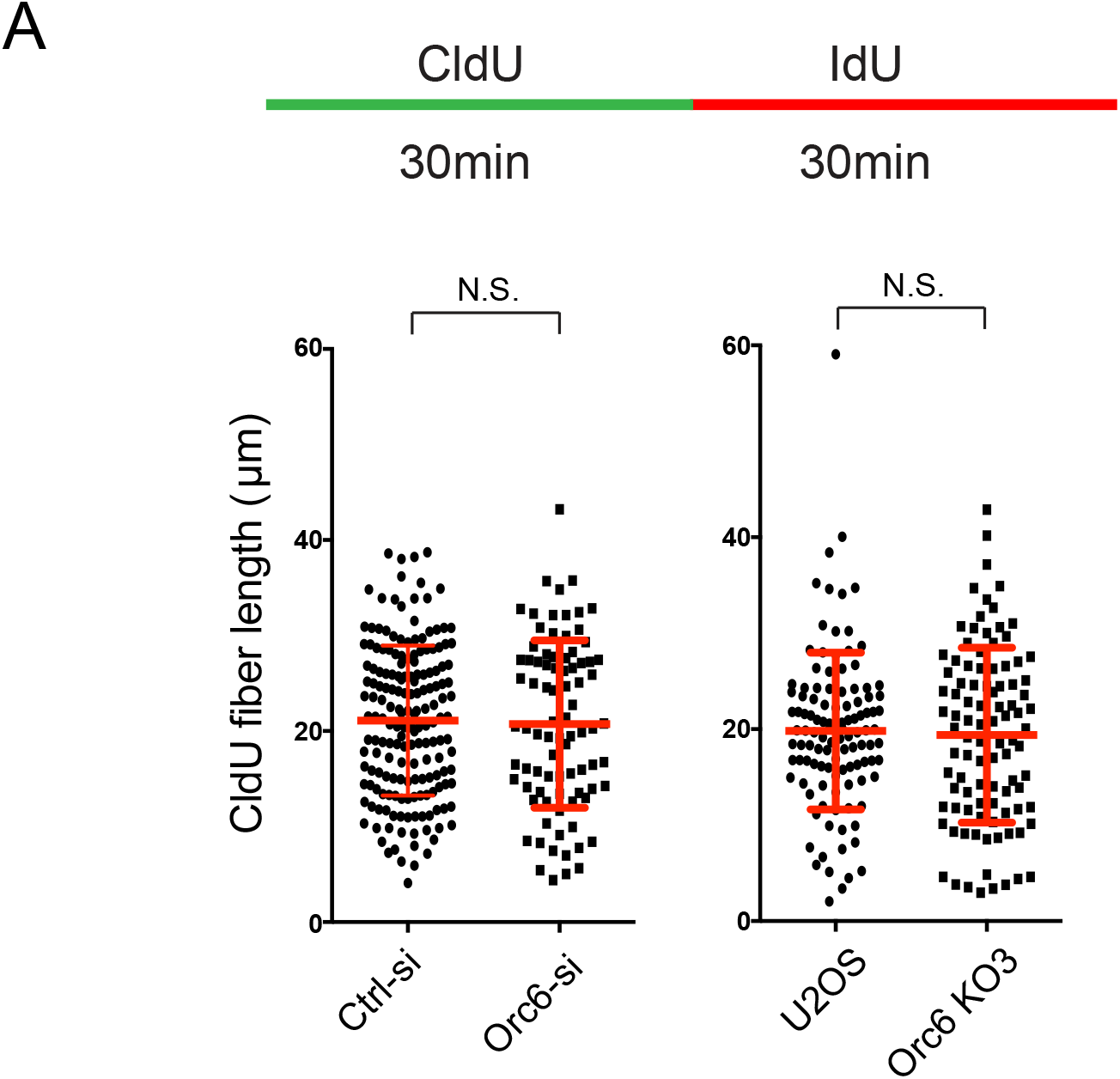
Loss of Orc6 does not alter replication fork rate (Related to figure 3) (A) DNA fiber assay was used to monitor replication fork progression speed. Mean ± SD.

**Figure S4.**
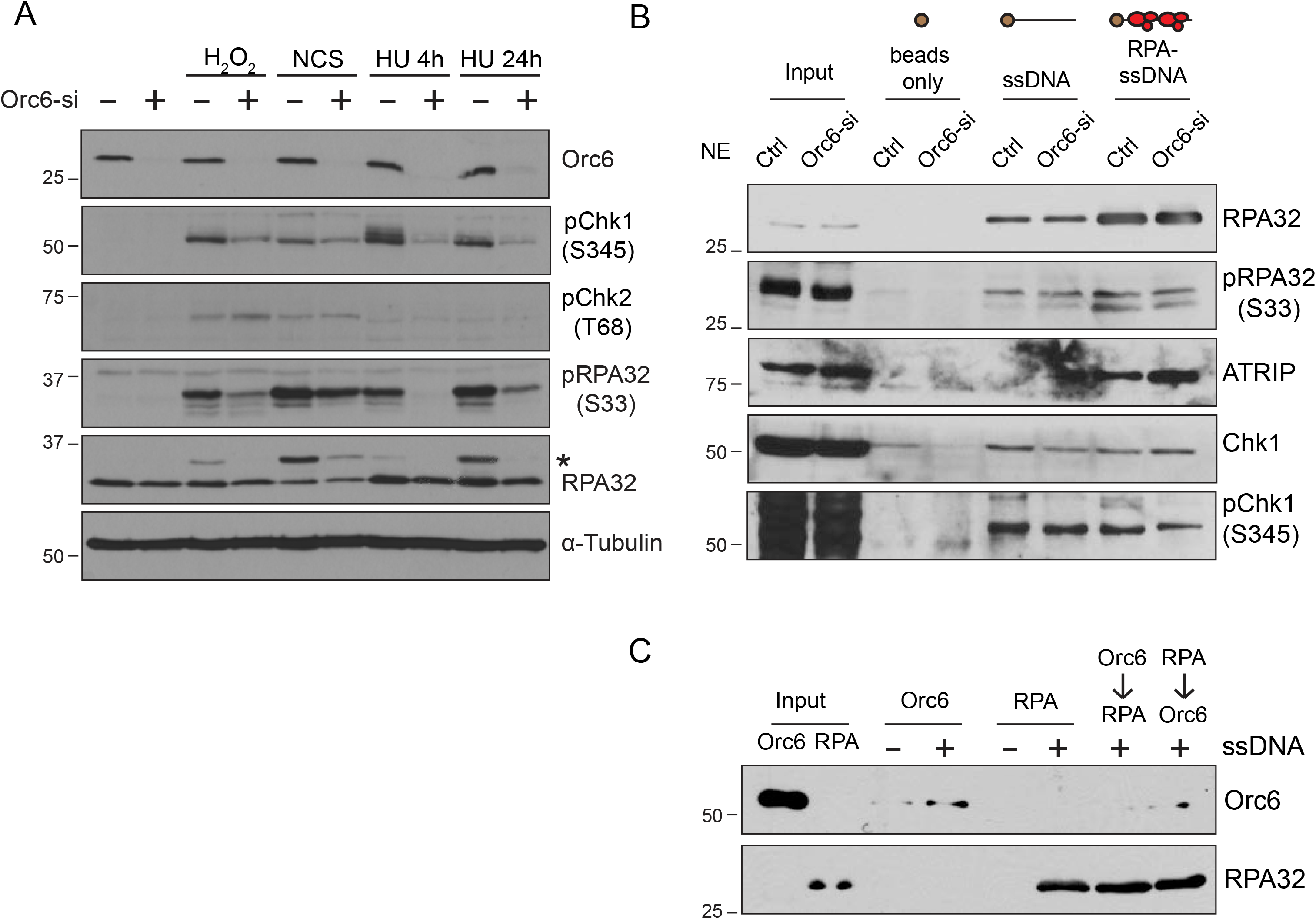
Reduced ATR activation in Orc6-depleted cells is not due to changes in ATR signaling proteins’ recruitment to RPA-ssDNA (Related to figure 5) (A) Western blot of ATM/ATR signaling pathways for different DNA damage drug treatment in control and Orc6 knockdown cells. *Asterisk* indicates hyperphosphorylated RPA32. (B) In vitro ssDNA pull-down assay to determine the recruitment of ATR signaling proteins. ssDNA along or ssDNA pre-coated with purified RPA were incubated with control or Orc6-depleted nuclear extracts. Samples obtained after biotin-ssDNA pull-down were analyzed by western blotting. NE, nuclear extract. (C) In vitro ssDNA pull-down for effects of Orc6 to RPA’s ssDNA binding ability. Orc6 along, RPA along, Orc6 first then RPA or RPA first then Orc6 were added to ssDNA. Proteins bound to ssDNA were analyzed using western blot.

**Figure S5.**
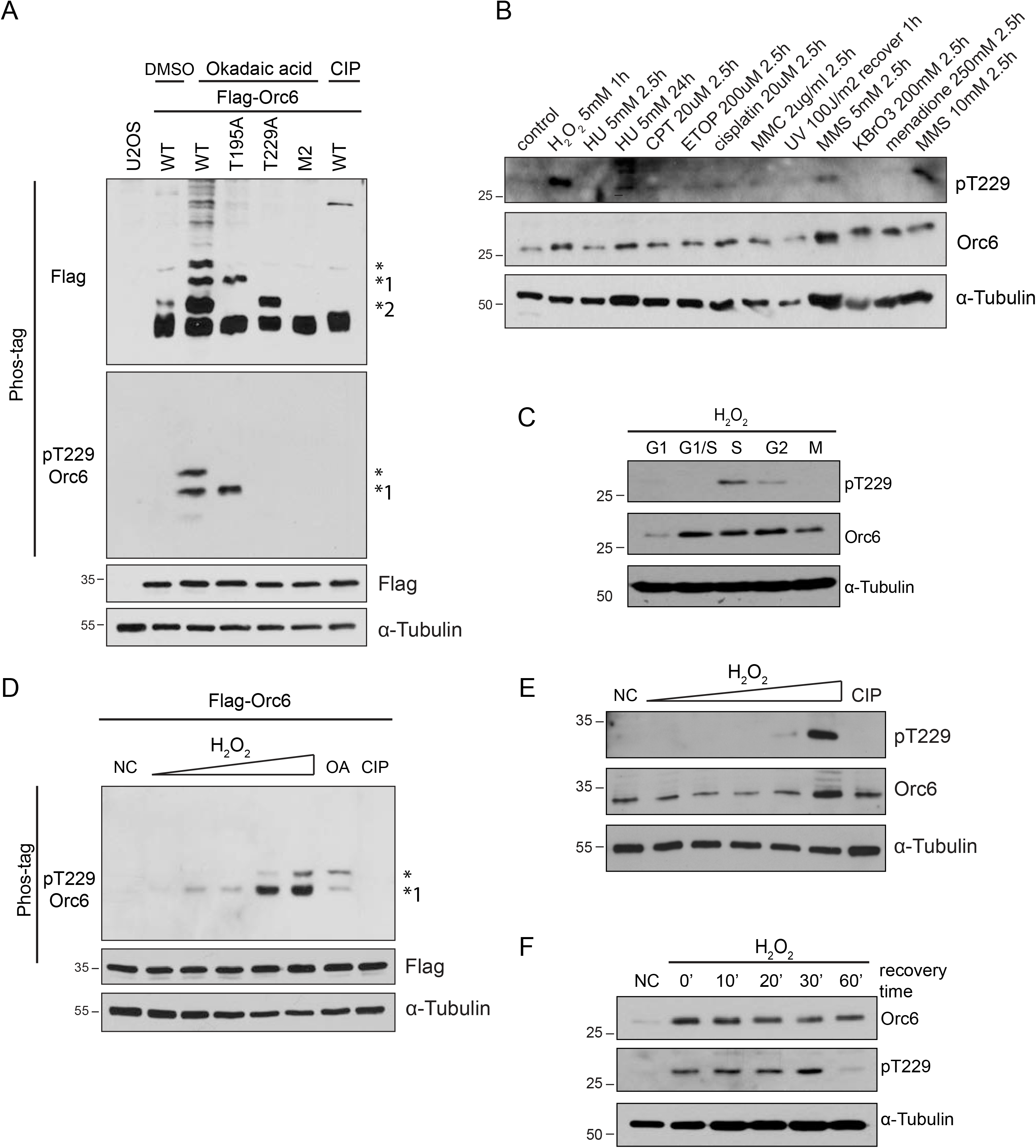
Orc6 is phosphorylated at Thr229 upon oxidative DNA damage (Related to figure 5) (A) Phos-tag gel analysis of Orc6 phosphorylation. U2OS cells were transfected with Flag-Orc6-WT, T195A, T229A or M2 (both T195 and T229 were mutated to Ala), and treated with okadaic acid to induce accumulation of phosphorylation. * corresponds to Orc6 phosphorylated on both T195 and T229; *1 indicates T229 phosphorylation. *2 indicates T195 phosphorylation. CIP, calf intestinal phosphatase. (B) Western blot for testing Orc6 T229 phosphorylation upon different genotoxic drug treatments. (C) Western blot showing Orc6 T229 phosphorylation pattern during cell cycle. (D) Phos-tag gel analysis to determine the dose-dependent effects of T229 phosphorylation upon H_2_O_2_ treatment. U2OS cells were transfected with Flag-Orc6-WT and treated with different concentration of H_2_O_2_ (0.25, 0.5, 1, 2, 5 mM). OA, okadaic acid. (E) Western blot for endogenous Orc6 T229 phosphorylation. (F) Western blot of T229 phosphorylation regulation. Cells were collected at indicated time point after release from 20 min H_2_O_2_ treatment. NC, negative control.

**Figure S6.**
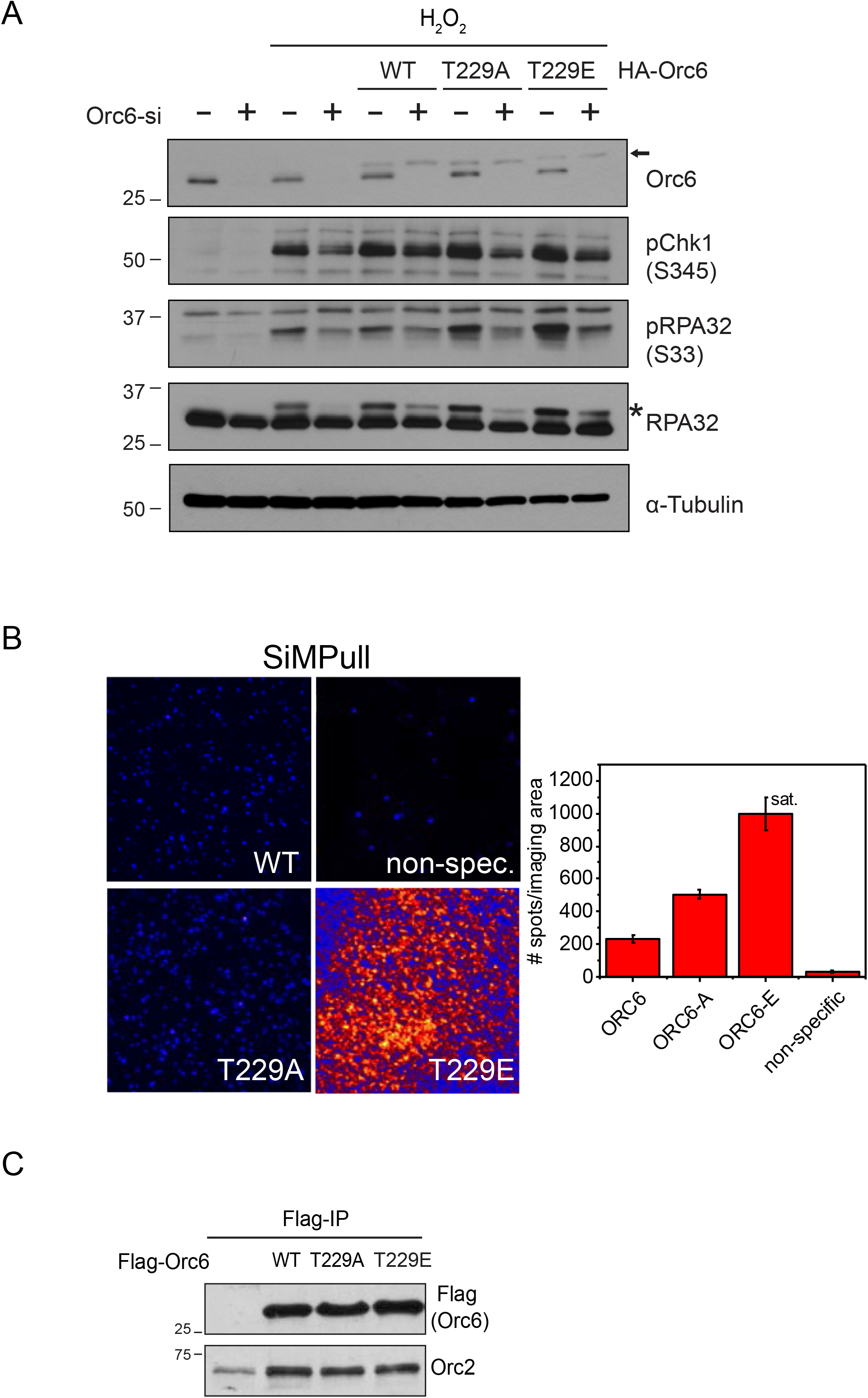
Analysis of Orc6 T229 mutants (Related to figure 5) (A) ATR activation analyzed by western blot of various U2OS cells (HA-Orc6-WT, HA-Orc6-T229A, HA-Orc6-T229E) depleted of endogenous Orc6. *Arrow* indicates HA-Orc6. *Asterisk* indicates hyperphosphorylated RPA32. (B) SiMPull for investigating the effect of Orc6 T229 phosphorylation to its DNA fork structure binding ability. Representative images of different mutants of purified GST-Flag-Orc6 (left) and quantified results (right). Mean ± SD. (C) Immunoprecipitation of different mutants of Flag-Orc6 from U2OS cells. Samples were analyzed by western blotting.

**Figure S7.**
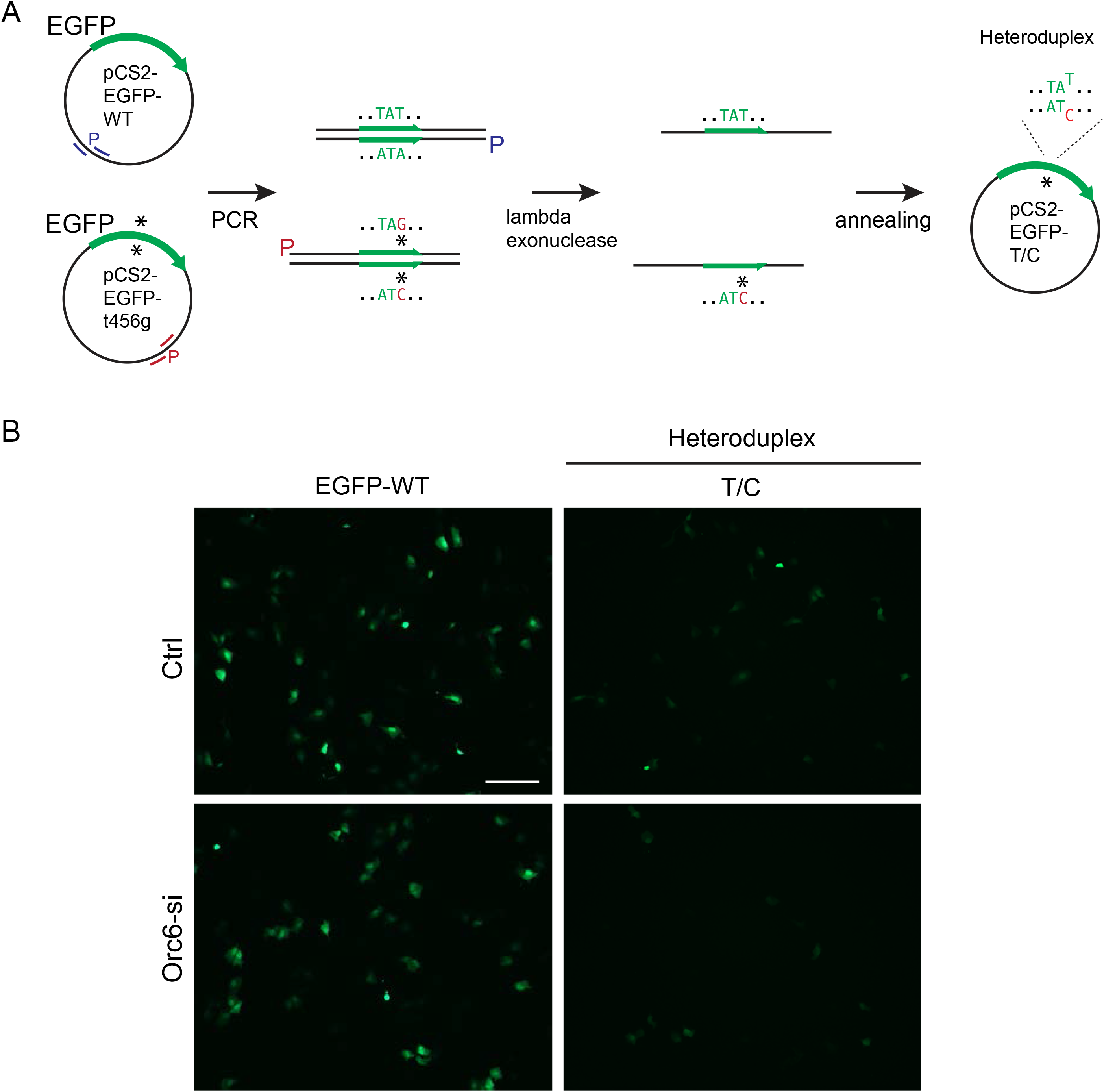
MMR in vivo reporter assay (Related to figure 7) (A) Schematic illustration of the generation of the heteroduplex used for reporter assay. (B) Representative images of MMR assay are shown. Scale bar, 200µm.

**Table S1. Number of peptides, sum intensities and relative values of proteins identified by mass spectrometry in control, T7-Orc6, and T7-Orc6 with H_2_O_2_ treatment**

**Table S2.**
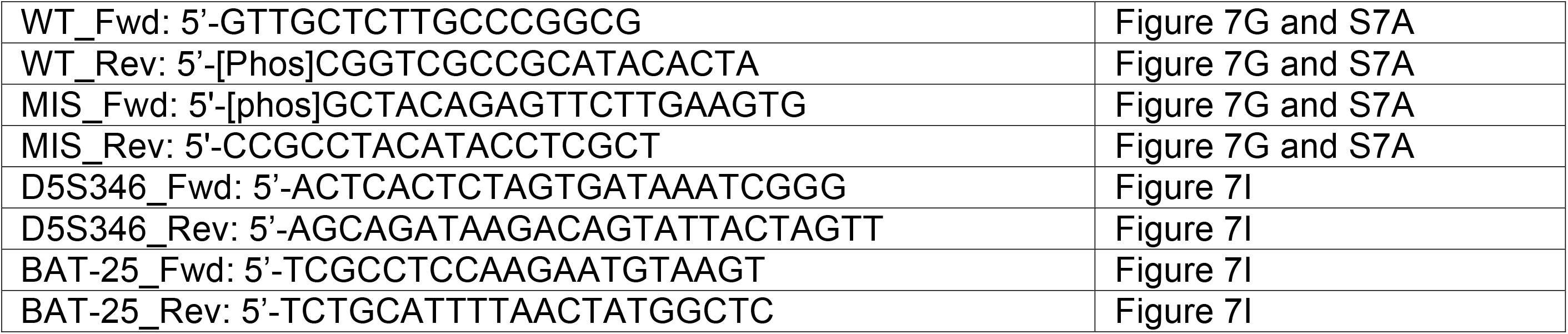
Primers used in this study.

## Notes

### Competing Interest Statement

The authors have declared no competing interest.

